# Imetelstat-Mediated Alterations in Fatty Acid Metabolism To Induce Ferroptosis As Therapeutic Strategy for Acute Myeloid Leukemia

**DOI:** 10.1101/2023.04.25.538357

**Authors:** Claudia Bruedigam, Amy H. Porter, Axia Song, Gerjanne Vroeg in de Wei, Thomas Stoll, Jasmin Straube, Leanne Cooper, Guidan Cheng, Vivian F. S. Kahl, Alexander P. Sobinoff, Victoria Y. Ling, Billy Michael Chelliah Jebaraj, Yashaswini Janardhanan, Rohit Haldar, Laura J. Bray, Lars Bullinger, Florian H. Heidel, Glen A. Kennedy, Michelle M. Hill, Hilda A. Pickett, Omar Abdel-Wahab, Gunter Hartel, Steven W. Lane

## Abstract

Telomerase enables replicative immortality in most cancers including acute myeloid leukemia (AML). Imetelstat is a first-in-class telomerase inhibitor with clinical efficacy in myelofibrosis and myelodysplastic syndromes. Here, we develop an AML patient-derived xenograft (PDX) resource, and perform integrated genomics, transcriptomics, and lipidomics analyses combined with functional genetics to identify key mediators of imetelstat efficacy. In a randomized Phase II-like preclinical trial in PDX, imetelstat effectively diminishes AML burden, and preferentially targets subgroups containing mutant *NRAS* and oxidative stress-associated gene expression signatures. Unbiased, genome-wide CRISPR/Cas9 editing identifies ferroptosis regulators as key mediators of imetelstat efficacy. Imetelstat promotes the formation of polyunsaturated fatty acid-containing phospholipids, causing excessive levels of lipid peroxidation and oxidative stress. Pharmacological inhibition of ferroptosis diminishes imetelstat efficacy. We leverage these mechanistic insights to develop an optimized therapeutic strategy using oxidative stress-inducing chemotherapy to sensitize patient samples to imetelstat causing significant disease control in AML.

## INTRODUCTION

Acute myeloid leukemia (AML) is an aggressive and lethal blood cancer with a 5- year overall survival rate of less than 45% for patients younger than 60 years of age, and less than 10% for older patients, predominantly due to disease relapse after chemotherapy or targeted treatments ^1^. AML has been extensively classified based on biological features, and advances in sequencing technologies have led to a comprehensive genetic classification strategy (European LeukemiaNet, ELN2017) ^2, 3^. Despite this improved understanding of the individual disease subtypes, targeted treatment algorithms have resulted in only modest clinical benefits to date ^4^. The development of effective therapies to improve remission rates and prevent relapse remains a top priority for patients with AML.

Telomerase is an attractive target as it is highly expressed and reactivated in the majority of AML, and absent in most cell types including normal hematopoietic cells ^5–7^. We have previously shown that genetic depletion of telomerase eradicates leukemia stem cells, particularly upon enforced replication ^8^. Despite promising preclinical evidence, the development of effective and specific telomerase inhibitors has been challenging. Imetelstat is a first-in-class covalently-lipidated 13-mer thiophosphoramidate oligonucleotide that can competitively inhibit telomerase activity by binding to the telomerase RNA component TERC ^9^. Imetelstat has shown clinical efficacy in essential thrombocythemia ^10^, myelofibrosis ^11^, and lower risk myelodysplastic syndromes ^12^. In myelodysplastic syndromes, clinical benefits are associated with reductions in telomerase activity and TERT expression ^12^.

In addition to its canonical role as critical regulator of telomere length maintenance, telomerase fulfils important non-canonical roles contributing to stress elimination, regulation of Wnt/beta-catenin, NF-κB and p65 signalling, as well as resistance to ionizing radiation ^13^. Hence, the clinical activity of imetelstat may be driven by mechanisms independent of telomere shortening, and potentially canonical telomerase activity ^14^.

Preclinical trials in patient-derived xenografts (PDX) provide genetically diverse, tractable models to define the efficacy of drugs and to identify biomarkers of response and resistance in AML ^15^. PDX-based trials also allow, within the same cohort, the evaluation of novel combination therapies with agents that may enhance efficacy, and also critically compare their additive value to current, established standard treatments.

In this study, we aimed to assess the preclinical efficacy of imetelstat in a large AML PDX resource that reflects the diversity of genetic abnormalities found in large patient cohorts. We utilized this AML PDX resource to identify biomarkers of resistance and response to imetelstat therapy, and to test potentially synergistic combination therapies. To elucidate the mechanism of action of imetelstat in an unbiased manner, we performed genome-wide CRISPR/Cas9 editing allowing the identification of gene knockouts that confer resistance to imetelstat therapy. This study reveals that imetelstat is a potent inducer of ferroptosis which effectively diminishes AML burden and delays relapse following oxidative stress inducing therapy.

## RESULTS

### Generation of a comprehensive and representative AML PDX resource

In order to generate a representative AML PDX inventory, primary bone marrow or blood samples from 50 patients were tested for engraftment and development of AML in NOD/SCID/IL2gR-/-/hIL3,CSF2,KITLG (NSGS) ^16^. The overall success rate for primary engraftment in NSGS was 70%, defined by bone marrow, spleen or peripheral blood donor chimerism of at least 20%, splenomegaly (spleen weight > 70 mg), anemia (HCT < 35%) or thrombocytopenia (PLT < 400×10^6^/ml), microscopically visible AML infiltration into spleen or liver, and peripheral blood blast morphology (***Supplementary Figures 1A-N)***. Successfully engrafted NSGS recipients developed AML with a median onset of 173 days post-transplant (***Supplementary Figure 1P***).

From the individual AML patient samples that successfully engrafted in NSGS, 30 were randomly selected and characterized based on clinical parameters including patient age, gender, ELN2017 risk, WHO disease classification, and molecular profiles obtained by transcriptional and mutational sequencing (***Figures 1A-C***). All ELN2017 prognostic risk (favorable, intermediate, adverse) and age categories were represented, 17 samples were from female, and 13 samples from male AML patient donors (***Figure 1B***). Oncogenic mutations were most frequently detected in *NPM1*, *DNMT3A*, and *FLT3* loci, and overall, this AML PDX resource recapitulated the genetic abnormalities that are observed in large clinical AML cohorts ^3^ (***Figure 1C***).

**Figure 1:**
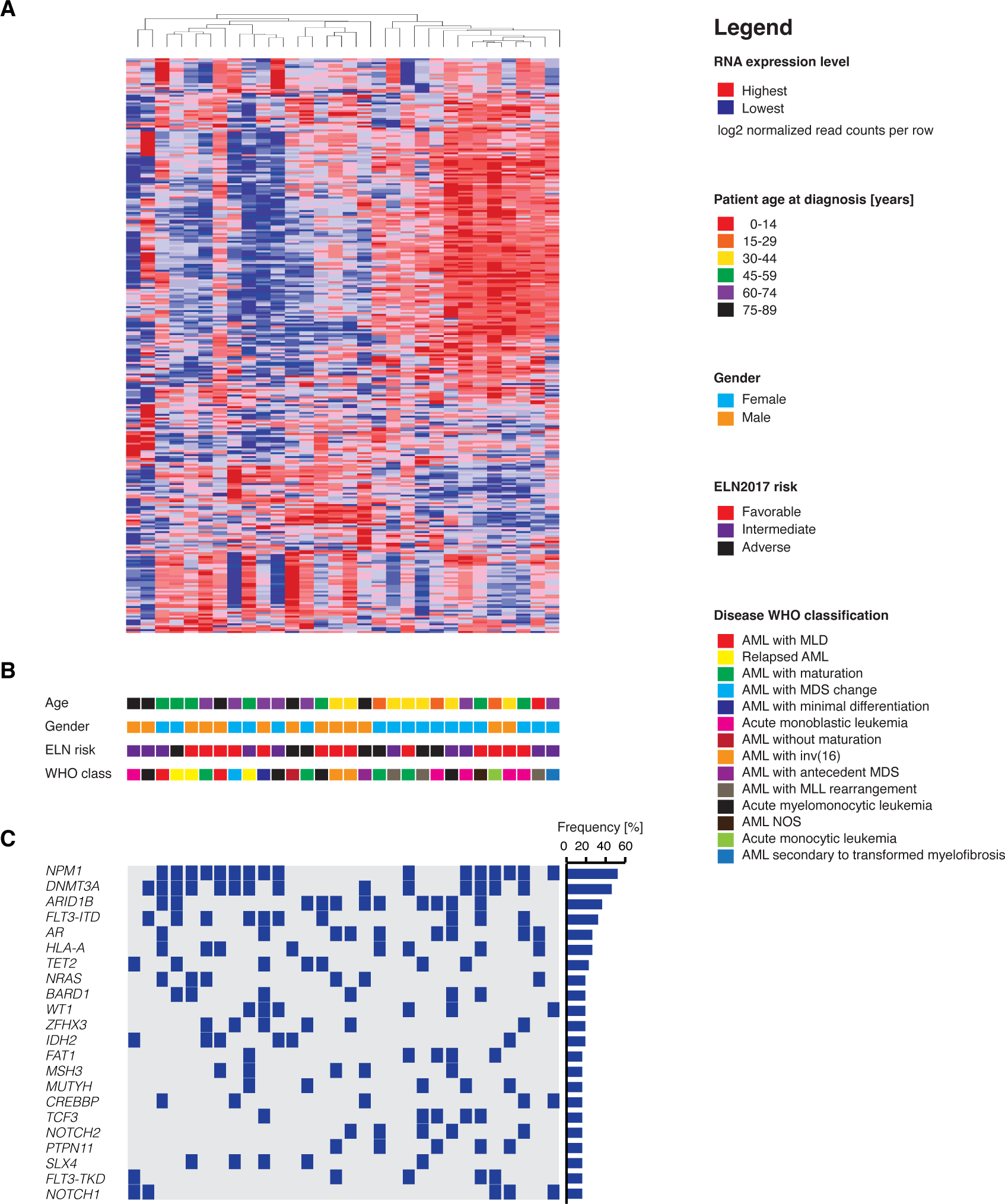
Integrative Analysis of AML Patient Samples. (A) Unsupervised hierarchical clustering analysis on the expression of 300 transcripts with the greatest variance-to-mean ratios among 30 individual AMLs from our repository that can successfully generate AML PDX. B) Key clinical characteristics of patients from whom AML samples were derived including age at diagnosis, gender, ELN2017 prognostic risk group, and WHO class of disease. C) OncoPrint of the most frequently detected mutations in AMLs by targeted next generation sequencing of 585 genes associated with hematologic malignancies (the MSKCC HemePACT assay) ^40^.

### Imetelstat diminishes AML burden and prolongs survival in a randomized Phase II-like preclinical trial in AML PDX

In order to test the pre-clinical efficacy of imetelstat in AML, the characterized 30 individual AML patient samples were each transplanted into 12 NSGS recipients (n = 360 PDX in total). Once AML burden was detected, PDX were randomized and treated with imetelstat or vehicle control (PBS) until disease onset or a survival benefit of at least 30 days was reached. Median survival was significantly prolonged in imetelstat compared to PBS-treated PDX (155 vs. 100 days post-start of treatment, p < 0.0001; ***Figure 2A***). AML burden measured as peripheral blood donor chimerism per day was significantly lower in imetelstat compared to vehicle-treated recipients (***Figure 2B***). Moreover, endpoint peripheral blood donor chimerism, bone marrow cellularity and donor chimerism as well as the absolute number of AML patient-derived cells were significantly reduced in recipients treated with imetelstat when compared to vehicle control (***Figures 2C-F***). Furthermore, imetelstat treatment significantly reduced splenic AML donor chimerism (***Figure 2G***). We next assessed AML surface marker expression associated with leukemia initiating activity ^17–19^ (***Figure 2H***). Imetelstat significantly diminished the CD34+CD38- LSC-enriched splenic AML cell population (***Figure 2I***). In normal human hematopoiesis using two independent CD34-enriched cord blood xenografts in NSG recipients, the effects of imetelstat were predominantly seen in B-lymphocytes with relative preservation of the myeloid and stem cell population (***Supplementary Figure 2***).

**Figure 2:**
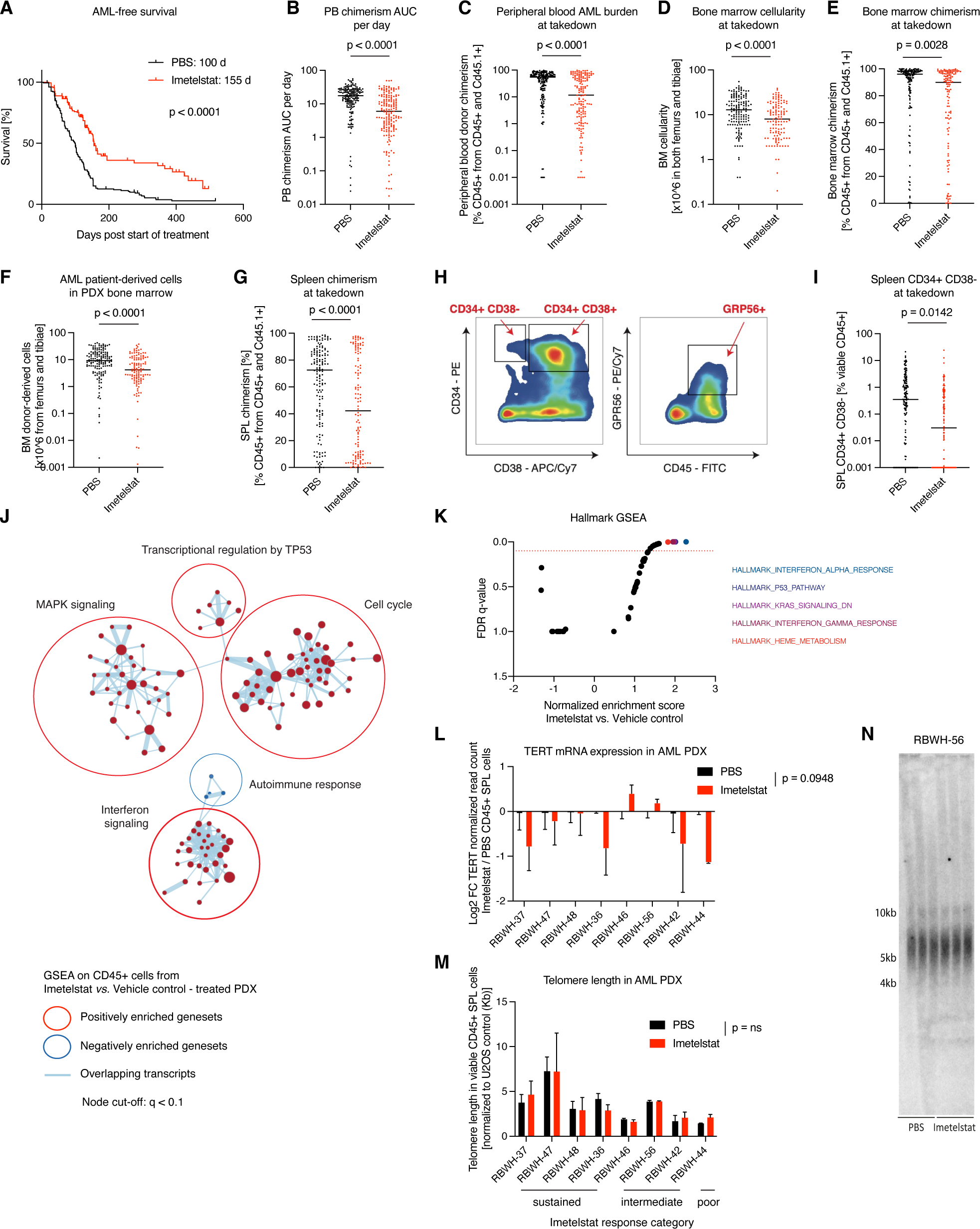
The efficacy of imetelstat in a randomized Phase II - like preclinical trial in AML PDX. Analysis of AML PDX that received imetelstat therapy intraperitoneally three times per week at 15 mg/kg body weight, or vehicle control (PBS) continuously until individual disease onset or a survival benefit of at least 30 days was reached. Thirty individual AML patient samples were transplanted into each twelve NSGS recipients, of which six were treated with imetelstat, and six were treated with PBS. PDX generated from the thirty individual AMLs were pooled into imetelstat-treated or PBS-treated groups with n = 180 per group. (A) Median AML-free survival was 100 (PBS) and 155 (imetelstat) days from start of treatment; p < 0.0001 according to Gehan-Breslow-Wilcoxon; n = 180 per group. (B-G) Quantification of peripheral blood donor chimerism area under the curve (AUC) per day (B), endpoint peripheral blood donor chimerism (C), bone marrow cellularity (D), bone marrow chimerism (E), the number of AML donor-derived cells in PDX bone marrow (F), and splenic donor chimerism (G). (H-I) Flow cytometric analysis of AML surface marker expression CD34, CD38 and GPR56. Gating strategy (H). Quantification of CD34+ CD38- viable CD45+ splenic singlets harvested from imetelstat or PBS-treated AML PDX (I). Statistical analysis was based on two-sided Student’s t test; n = 180 PDX per group. Solid lines represent the mean of each group. (J-K) RNAseq analysis on imetelstat or vehicle control-treated AML PDX. Gene set enrichment analysis (GSEA) on RNAseq data from sorted hCD45+ cells harvested from imetelstat or vehicle control (PBS) - treated AML PDX. N = 8 individual AML patient samples with n = 2 PDX per AML patient sample and treatment group. Cytoscape nodes represent gene sets with a cut-off: q < 0.1 (J); GSEA focused on hallmark signatures with the top five enriched signatures highlighted in color (K). (L-N) Analysis of the effect of imetelstat treatment on telomerase / telomere status. TERT mRNA expression results obtained from RNAseq analysis described as above (L). Telomere length in viable CD45+ splenic cells from imetelstat versus PBS-treated AML PDX measured by qPCR (M) and confirmed by telomeric restriction fragment analysis (N). Statistical analysis (L-M) was based on paired Student’s t-test comparing imetelstat with vehicle groups within the respective AML patient samples.

We next aimed to compare imetelstat responses to those obtained with standard induction chemotherapy (i.e. cytarabine plus anthracycline) in AML PDX from 20 individual AML patient samples in an independent cohort using NOD.Rag1-/-Il2Rg-/- / hIL3,CSF2,KITLG (NRGS) recipients ^20^. Imetelstat matched the similar benefit conveyed by standard chemotherapy (139 days) comparative to 104 days in the vehicle control group, and this was accompanied by significant reductions in peripheral blood AML burden (***Supplementary Figures 3A-B***). However, the individual AML patient samples could be classified into either preferential imetelstat or preferential chemotherapy responders (***Supplementary Figure 3C***). Preferential responses to imetelstat when compared to standard induction chemotherapy were associated with baseline mutations in *NRAS*, *JAK2*, or *GLI1* (***Supplementary Figure 3D***).

We next assessed the transcriptional consequences of imetelstat therapy in a cohort of PDX from eight randomly chosen individual AML patient samples *in vivo* (n = 4 sustained (RBWH-37, -47, -48, -36), n = 3 intermediate (RBWH-46, -56, -42), and n = 1 poor (RBWH-44) responders to imetelstat). Gene expression signatures annotated as interferon signaling, cell cycle, transcriptional regulation by TP53, and MAPK signaling were significantly enriched in AML donor cells from imetelstat-treated compared to vehicle control-treated PDX (***Figures 2J-K***). TERT mRNA expression levels were trend-wise reduced in AML donor cells derived from imetelstat-treated compared to vehicle-treated PDX spleens (***Figure 2L***). Intriguingly, telomere lengths were similar between imetelstat-treated compared to vehicle-treated groups (***Figures 2M-N***).

### Identification of key mediators of imetelstat efficacy using genome-wide CRISPR/Cas9 editing

In order to investigate the mechanism of action of imetelstat in AML in an unbiased manner, we applied the Brunello sgRNA library ^21, 22^ as positive selection screen to identify gene knockouts that confer resistance to imetelstat. We used NB4 cells as these demonstrated highest sensitivity to imetelstat when compared to thirteen other human hematopoietic cell lines (***Supplementary Figure 7A***). IC50 values strongly depended on cell density, demonstrating the presence of an imetelstat inoculum effect (***Supplementary Figure 4A)*** ^23, 24^. Cas9 expressing NB4 cells transduced with the Brunello library or untransduced controls were cultured in the presence of imetelstat concentrations that resulted in significant cell death of the untransduced control cultures but allowed the enrichment of imetelstat-resistant cells in Brunello-transduced cultures over a timecourse of 45 days in culture (***Supplementary Figure 4B***). Vehicle or mismatch control treated NB4 cells grew exponentially throughout the course of treatment (***Supplementary Figure 4C***). Specific guide RNAs were selectively enriched in Brunello-transduced imetelstat-resistant compared to vehicle-treated and input control cultures (***Supplementary Figures 4D-G***). Combined RIGER and STARS gene-ranking algorithms identified seven significant hits: fatty acid desaturase 2 (*FADS2)*, acyl-CoA synthetase long chain family member 4 (*ACSL4)*, translocase of inner mitochondrial membrane 17A (*TIMM17A)*, late endosomal/lysosomal adaptor, MAPK and MTOR activator 1-3 (*LAMTOR1, LAMTOR2*, *LAMTOR3*), and myosin regulatory light chain interacting protein (*MYLIP;* ***Figure 3A***). Ingenuity pathway analysis indicated close functional relationships between the seven hits in regulating lipid metabolism, iron / metal ion binding, mitochondrial matrix, and lysosome biogenesis and localization (***Figure 3B***).

**Figure 3:**
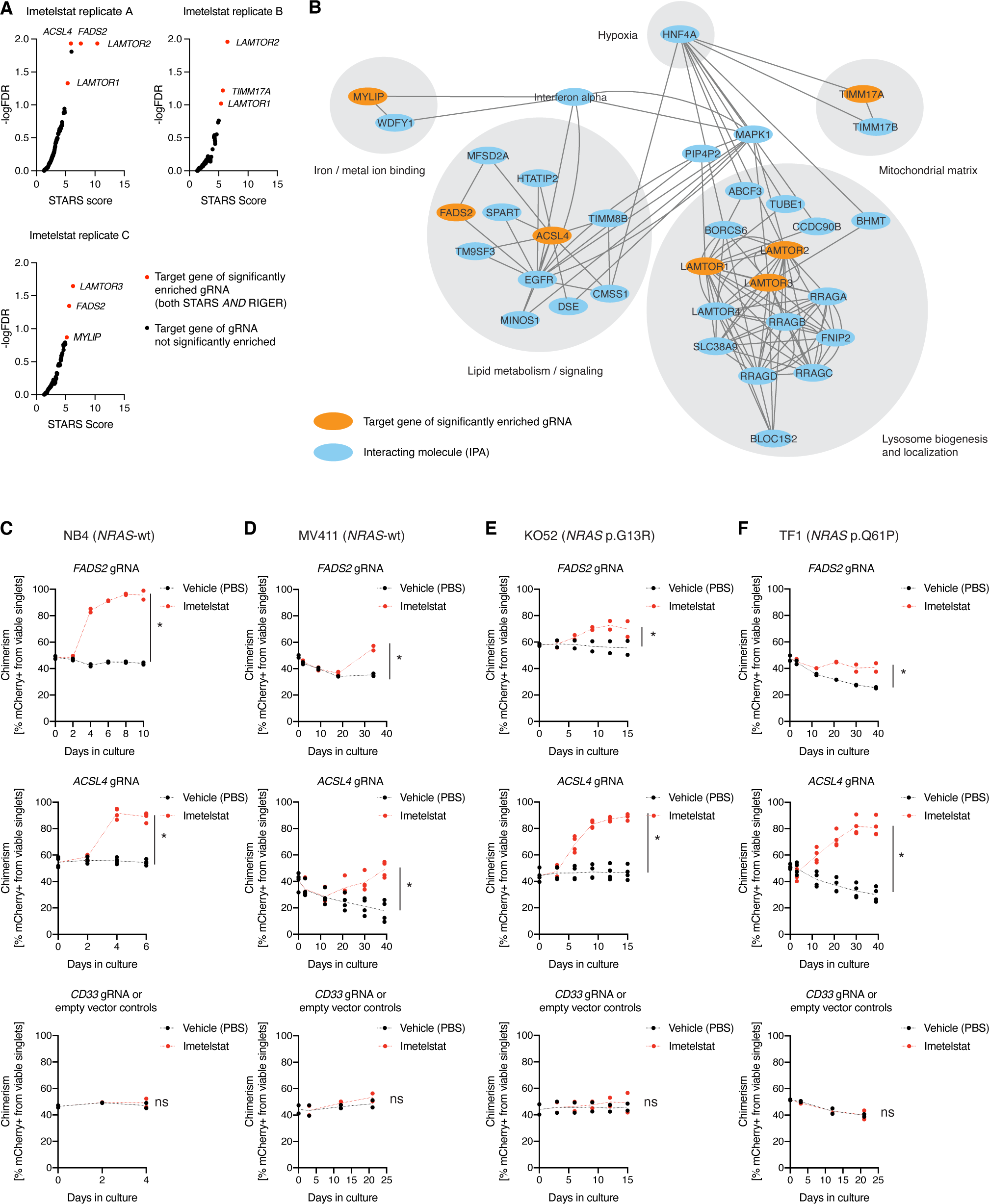
Identification of key mediators of imetelstat efficacy using genome- wide CRISPR/Cas9 editing. Brunello CRISPR/Cas9 positive enrichment screen in NB4 cells. Cas9-expressing cells were generated to contain one single guide RNA from this library each cell. These cultures and untransduced control cultures were grown in the presence of vehicle control (PBS), mismatch control (MM1), or imetelstat for 45 days, allowing the growth of an imetelstat-resistant culture with an IC98 that corresponded to approximately 45 cell doublings. (A) Guide RNA enrichment analysis using STARS *AND* RIGER gene-ranking algorithms in the three independent imetelstat-treated biological replicates. Red circles indicate significantly enriched targets (STARS FDR < 0.15 *AND* RIGER score > 2.0). (B) Cytoscape visualization of the IPA-derived interaction network connecting the identified significantly enriched guide RNA targets. (C-F) Competition assays of imetelstat (red) *versus* vehicle control (PBS; black) treated Cas9-expressing NB4 (C), MV411 (D), KO52 (E), and TF1 (F) cultures transduced with two independent single guide RNAs targeting *FADS2* (top panel), four independent single gRNAs targeting *ACSL4* (middle panel), and empty vector control as well as two independent guide RNA constructs targeting *CD33*. At least 2 independent experiments from separate cell passages with n = 3 replicates per condition were pooled. Asterisks (*) denote statistically significant differences (95% confidence interval) between chimerism AUC from imetelstat vs. PBS-treated cell cultures.

We next aimed to validate the most significant hits identified (i.e. *FADS2* and *ACSL4*) using single guide RNA-mediated editing in the *NRAS*-wild type expressing NB4 and MV411, and the *NRAS*-mutant KO52 (p.G13R) and TF1 (p.Q61P) AML cell lines. Editing was confirmed by TIDE analysis ^25^ and reduced protein levels (***Supplementary Figures 4H-J***).

We performed competition assays to confirm that loss-of-function editing of *FADS2* or *ACSL4* confers competitive growth advantage under imetelstat pressure in all AML cell lines analyzed (***Figures 3C-F***). The observed effects were target-specific as a competitive outgrowth under imetelstat pressure was not observed when CD33 (predicted to have neutral effects on cell functions ^26^) knockouts or empty vector controls were used (***Figures 3C-F, bottom panel***).

These results demonstrate that loss-of-function editing of FADS2 or ACSL4 confers competitive growth advantage under imetelstat pressure, identifying ACSL4 and FADS2 as mediators of imetelstat efficacy in AML.

### Imetelstat is a potent inducer of ferroptosis

*ACSL4* and *FADS2* encode key enzymes regulating polyunsaturated fatty acid (PUFA)-containing phospholipid synthesis. FADS2 is a key enzyme in a lipid metabolic pathway that converts the essential fatty acids linoleate (18:2n6) and α- linolenate (C18:3n3) into long-chain PUFAs ^27^. Targeted lipidomics analysis on 593 lipid species and their desaturation levels ^28, 29^ demonstrated clear effects of imetelstat treatment and *FADS2* editing on the cellular lipidome, with imetelstat-treated empty vector control AML cells showing greatest difference to vehicle treated empty vector control and *FADS2*-edited cells. (***Supplementary Figure 5A***). Imetelstat increased the levels of phospholipids with triglycerides and reduced the levels of phospholipids containing cholesteryl esters and ceramides when compared to vehicle control in an FADS2-dependent manner (***Supplementary Figure 5C***). Using lipidr analysis package ^29^, we found a significant enrichment of phospholipids containing fatty acids with three unsaturated bonds in imetelstat-treated compared to vehicle control-treated NB4 cells, and this enrichment of lipid desaturation was diminished by *FADS2* editing (***Figure 4A, Supplementary Figure 5B***). These data demonstrate imetelstat - induced PUFA phospholipid synthesis in an FADS2-dependent manner.

ACSL4 has been previously identified as key regulator of ferroptosis ^30^. Ferroptosis is a form of cell death that is driven by an imbalance between the production of ROS during lipid peroxidation and the antioxidant system, and may involve autophagic processes depending on the trigger ^31^. A hallmark of ferroptosis is lipid peroxidation, the oxidation of polyunsaturated fatty acid (PUFA)-containing phospholipids that occurs via a free radical chain reaction mechanism ^31^. Cancer therapies can enhance ferroptosis sensitivity via lipid remodeling that increases levels of peroxidation-susceptible PUFA-containing phospholipids ^32^.

To test whether imetelstat induces lipid peroxidation, we treated various AML cell lines with C11-BODIPY, a fluorescent fatty acid probe that changes its emission spectrum from red to green upon oxidation. In all 4 AML cell lines tested, imetelstat treatment resulted in a significant increase in MFI of the oxidized fatty acid probe, demonstrating that imetelstat induces lipid peroxidation in AML cells *in vitro* (***Figure 4B***). We next assessed whether also ROS levels were affected by imetelstat. Using Cellrox Green to measure ROS production, we found that its MFI was increased by imetelstat, and this increase was diminished when the lipid ROS scavenger ferrostatin-1 was added during the incubation step with cellrox green, demonstrating that imetelstat increases predominantly lipid ROS levels in AML cell lines *in vitro (****Figure 4C***). Both lipid peroxidation and lipid ROS production were significantly diminished in *ACSL4* or *FADS2* loss-of function edited AML cell lines, demonstrating that imetelstat-induced lipid peroxidation and lipid ROS production are dependent on functional FADS2 and ACSL4 *in vitro (****Figure 4D***; ***Supplementary Figure 6A***). Pharmacological inhibition of ferroptosis using the lipid ROS scavengers ferrostatin-1 and liproxstatin-1 diminished imetelstat efficacy in all AML cell lines tested (***Supplementary Figure 6B***). Moreover, the iron chelator DFOM, the 5- lipoxygenase inhibitor zileuton, and menadione diminished imetelstat-induced cell death in a significant proportion of AML cell lines tested (***Supplementary Figure 6B***).

**Figure 4:**
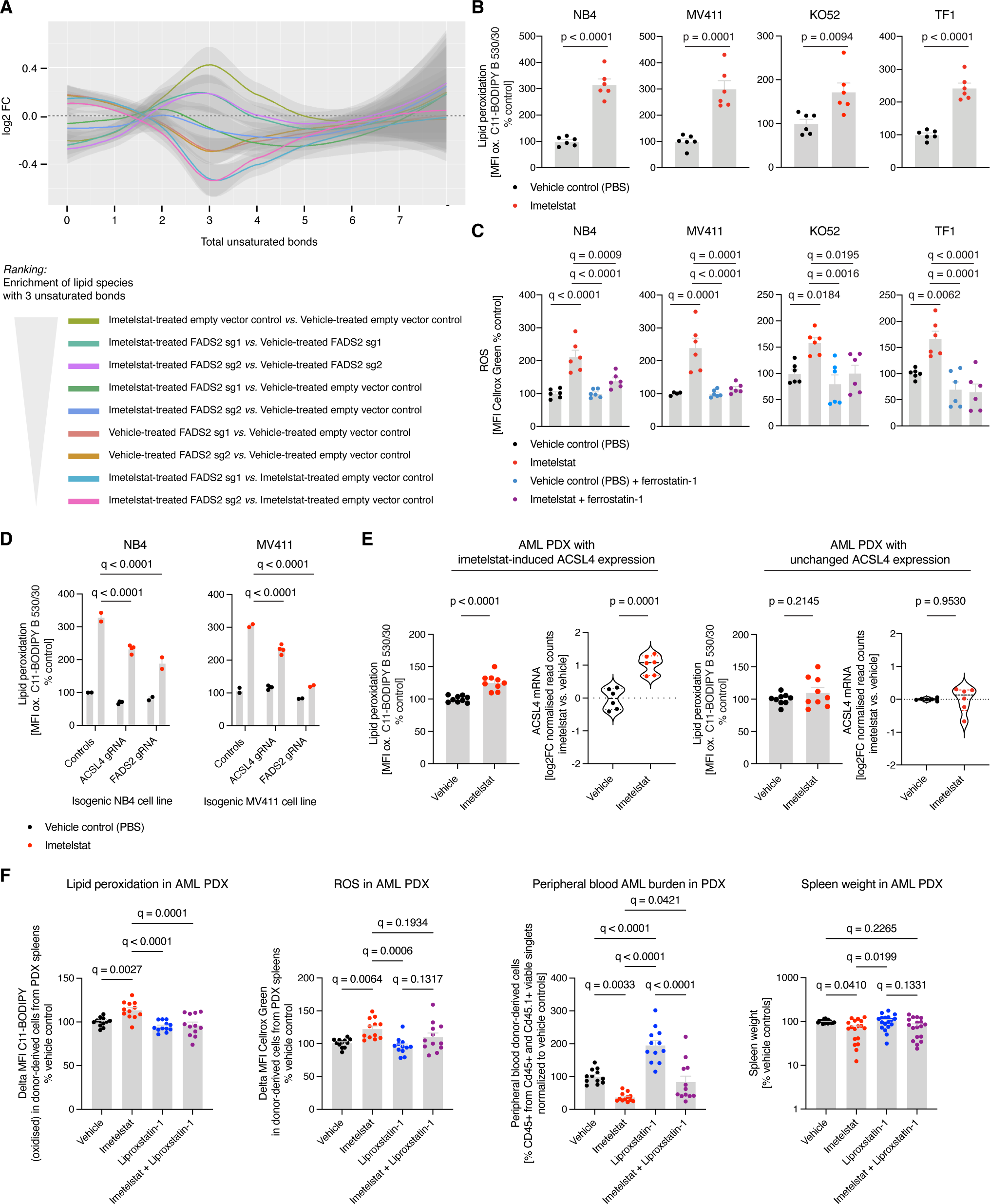
Imetelstat is a potent inducer of ferroptosis. (A) Lipid desaturation analysis of *FADS2*-edited (i.e. FADS2 sg1, FADS2 sg2) or non-edited (i.e. empty vector control) NB4 cells treated with imetelstat (4 μM at a seeding density of 2.5x10^5 cells per ml culture) or vehicle control for 24h. The graph depicts the log2 fold-change (FC) of the number of total unsaturated bonds in lipid species in the respective comparisons outlined in the panel legend. Shading represents the 95% confidence interval. N = 3 replicates from distinct cell passages. Additional lipidomics analyses are provided as supplementary methods information. (B-C) Flow cytometric analysis of lipid peroxidation using C11-BODIPY (B) and ROS levels using Cellrox Green (C) in NB4, MV411, KO52 and TF1 cell lines treated with imetelstat (4 μM at a seeding density 1-5x10^5 per ml). Timepoints of analysis: NB4 – 24h; MV411 – day 4; KO52 – day 8; TF1 – day 5 of culture. Statistics for (B) according to unpaired Student’s t-test with n = 6 replicates per condition. The four groups in (C) were analysed using One-way ANOVA with multiple comparisons. (D) Lipid peroxidation analysis in *ACSL4*-edited (i.e. ACSL4- sg1, ACSL4-sg2, ACSL4-sg3, ACSL4-sg4) or *FADS2*-edited (i.e. FADS2-sg1, FADS2-sg2) or non-edited (i.e. Cas9, empty vector) NB4 or MV411 cells treated with imetelstat (4 μM) or vehicle control for 24h. Statistics according to One-way ANOVA with multiple comparisons. (E) Lipid peroxidation analysis on sorted viable CD45+ splenic cells from imetelstat- compared to vehicle (PBS) – treated AML PDX from the preclinical trial presented in Figure 2. Patient samples were segregated into two cohorts based on differential ACSL4 expression at mRNA level (derived from RNAseq data presented in Figure 2). N = 6 individual AML PDX models. Statistics based on Student’s t-test. (F) *In vivo* rescue experiment using liproxstatin-1. PDX were generated from three individual AML patient samples with n = 4-6 per group, and treated with vehicle control, liproxstatin-1 (15 mg/kg body weight twice daily), imetelstat (15 mg/kg body weight three times per week), or the combination of liproxstatin-1 with imetelstat for two weeks. Lipid peroxidation and ROS levels were determined in CD45+ singlets from PDX spleens, and AML burden was measured by peripheral blood AML donor chimerism and spleen weight at the end of the treatment period. Statistics based on One-way ANOVA using multiple comparisons as indicated. In all panels, p-values or adjusted p-values (i.e. q-values) are displayed.

In AML PDX *in vivo*, imetelstat-induced lipid peroxidation was associated with increased ACSL4 expression (***Figure 4E***). To investigate whether lipid ROS and lipid peroxidation are essential for imetelstat’s mechanism of action in AML PDX *in vivo*, we treated AML PDX with either vehicle control, imetelstat (15 mg/kg three times per week), liproxstatin-1 (15 mg/kg twice daily) or the combination of both imetelstat and liproxstatin-1 for two weeks. Imetelstat-driven lipid peroxidation and ROS production were prevented by liproxstatin treatment (***Figure 4F***). *In vivo* liproxstatin treatment diminished imetelstat efficacy in PDX as measured by peripheral blood AML burden and spleen weight (***Figure 4F***).

Taken together, these data provide evidence that imetelstat is a potent inducer of ferroptosis through ACSL4- and FADS2-mediated alterations in PUFA metabolism, excessive lipid peroxidation and oxidative stress.

### Integration of transcriptomics with functional genetics identifies lipid droplet-associated G-quadruplex binding proteins as imetelstat target candidates mediating lipophagy-induced ferroptosis

By integrating transcriptomics and functional genetics, we aimed to investigate the mechanism by which imetelstat induces ferroptosis. We performed an overlay of the *in vivo* AML PDX RNAseq data sets from imetelstat and vehicle treated mice with the Brunello library CRISPR/Cas9 knockout screen data (cut-off criteria: RNAseq adjusted p-value < 0.05 *AND* RIGER p < 0.05), and identified eleven imetelstat target candidates (***Figure 5A***). Two of them, VIM (vimentin) and LMNA (lamin A/C), that are part of a common regulatory module (***Figure 5A***), have recently been identified as telomeric G-quadruplex binding proteins ^33^.

**Figure 5:**
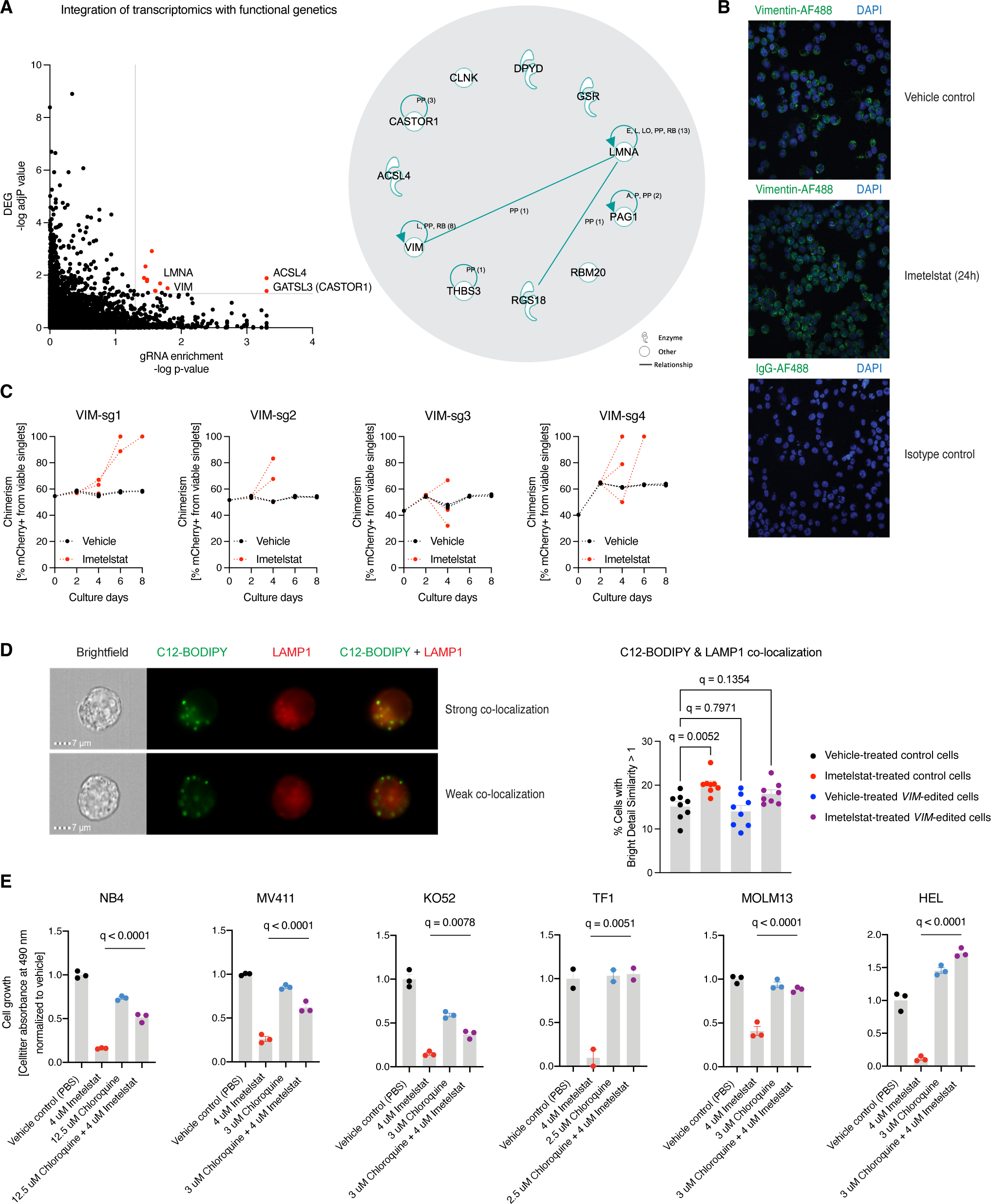
Integrative analysis of transcriptomics and functional genetics. (A) Integration of RNAseq and Brunello CRISPR screen data using relaxed cut-off criteria (DEG: adjusted p-value < 0.05 *AND* RIGER p-value < 0.05). Thirteen genes (red dots) passed these cut-off criteria, of which eleven were annotated in IPA (right panel). A common regulatory module for VIM, LMNA and RGS18 is highlighted through connecting lines. (B) Confocal microscopy to confirm VIM expression at protein level in NB4 cells treated with vehicle control or imetelstat for 24h. VIM protein levels appeared trend-wise increased but were not statistically different by confirmatory flow cytometry analysis (data not shown). (C) Loss-of-function editing of *VIM* in NB4 cells using four independent single gRNAs. Competition assays of mCherry+ *VIM*-edited cells grown in the presence of mCherry- unedited control NB4 cells, treated with imetelstat (red) or vehicle (PBS) control (black). (D) Imaging flow cytometry of lipophagy using the lipid droplet probe C12-BODIPY and late endosomal marker LAMP1 in *VIM*-edited (VIM sg1, VIM sg2, VIM sg3, and VIM sg4) or non-edited (i.e. native, Cas9, empty vector, CD33 sg2) NB4 cells. Recovery examples of cells showing strong co-localization of C12-BODIPY and LAMP1 indicative of lipophagy activity (top panel), or cells with weak co-localization indicating insignificant lipophagic flux. Quantification of the percentages of cells with strong co-localization defined as bright detail similarity score > 1. Statistics based on one-way ANOVA with multiple comparisons to vehicle treated control cells. Adjusted p-values (i.e. q-values) are displayed within the plot. (E) Chloroquine and imetelstat combination treatments in a panel of AML cell lines. Statistics based on one-way ANOVA with multiple comparisons. Significant q-values < 0.05 comparing imetelstat with imetelstat + chloroquine treated cultures are displayed in each graph.

Recent independent work demonstrated the capacity of imetelstat to form G-quadruplex structures *in vitro*, and this capacity is attributed to the presence of a triple G-repeat in its sequence ^34^. These insights prompted us to obtain an additional mismatch control harbouring a similar triple G repeat, but containing enough mismatches to prevent efficient binding to telomerase (***Supplementary Figure 7A***). Using an antibody raised against (T4G4)2 intermolecular G-quadruplex DNA structures ^35–37^, we found that imetelstat or GGG-containing mismatch but not mismatch 1 significantly interfered with endogenous G-quadruplex structures (***Supplementary Figure 7B***). In a panel of fourteen human hematopoietic cell lines, GGG-containing mismatch control and imetelstat demonstrated similar efficacies in the majority of AML cell lines tested (***Supplementary Figure 7A***). Moreover, GGG-containing mismatch was similarly effective as imetelstat in increasing ROS levels when compared to vehicle control (***Supplementary Figure 7C***). Ferrostatin- or DFOM- mediated inhibition of ferroptosis rescued both imetelstat as well as GGG- mismatch - induced cell death (***Supplementary Figure 7D***). We next compared the preclinical efficacy of imetelstat with GGG-mismatch, and mismatch 1 in an *NRAS/KRAS*-mutant AML PDX model (RCH-11). In this model, GGG-mismatch was also effective in reducing AML burden (***Supplementary Figure 7E***).

In addition to binding telomeric G-quadruplexes, vimentin has long been established as structural component of lipid droplets regulating their biogenesis and stability. Lipid droplets can undergo selective autophagy (i.e. lipophagy) that can result in the induction of ferroptosis ^38^. We hypothesized that imetelstat-induced PUFA-phospholipid synthesis, oxidation and ferroptosis can result from lipophagy. Vimentin was highly expressed at protein level in AML cells *in vitro* (***Figure 5B***), and loss-of-function editing of vimentin resulted in a modest competitive growth advantage of AML cells under imetelstat pressure (***Figure 5C***). We next assessed lipophagy using C12-BODIPY, a fluorescent fatty acid probe for lipid droplets, in conjunction with the late endosomal marker LAMP1^39^. Imaging flow cytometry revealed significantly increased co-localization of lipid droplets with the late endosomal marker LAMP1, indicating increased lipophagy (***Figure 5D***). To test whether pharmacological inhibition of lipophagy can prevent from imetelstat-induced ferroptosis, we cultured AML cells in the presence of imetelstat combined with chloroquine which inhibits lysosomal hydrolases by increasing the pH and thus lipophagy. Strikingly, in all AML cell lines tested, chloroquine diminished imetelstat-induced cell death (***Figure 5E***). These results provide evidence for a role of lipophagy-induced ferroptosis in imetelstat’s mechanism of action in AML via impaired lipid droplet homeostasis due to G-quadruplex mediated interference with the structural components of lipid droplets.

### Mutant *NRAS* and oxidative stress gene expression signatures associate with sustained responses to imetelstat

We next aimed to identify biomarkers of imetelstat response and resistance. Improved survival in imetelstat-treated AML PDX correlated with significantly reduced engraftment and disease burden, however there were clear differences in the magnitude and duration of individual responses (***Supplementary Figure 8***). To understand determinants of imetelstat response, we allocated each individual AML patient sample into either sustained, intermediate, or poor imetelstat response categories based on the individual effect of imetelstat on AML burden measured in peripheral blood over time (***Supplementary Figures 9D, 10A***). All ELN2017 prognostic risk categories were represented in each imetelstat response group, suggesting that the effects observed were not solely explained by favorable disease (***Supplementary Figure 10B***). In addition, cytogenetics, gender, age, FLT3-ITD allelic ratio, and TERT mRNA expression levels at baseline appeared similar among imetelstat response groups (***Supplementary Figures 10C-H***).

We next aimed to identify genetic biomarkers of response and resistance to imetelstat therapy by analyzing the data from individual AML patient samples at baseline that were generated by genomic sequencing using a comprehensive panel of 585 genes frequently mutated in hematological malignancies ^40^ (***Supplementary Figure 9C***). Oncogenic mutations in genes annotated in signaling or cell adhesion / metabolism were trend-wise more frequently observed in sustained compared to poor responders to imetelstat (***Figure 6A, Supplementary Figure 6C***).

Mutant *NRAS* was associated with enhanced responses to imetelstat therapy. This was evidenced by reduced AML burden and borderline-significant improvement in survival when compared to wild-type *NRAS* containing AML PDX (***Figures 6A-C***). Moreover, variant allelic frequencies of the relevant *NRAS* mutations inversely correlated with AML burden in imetelstat-treated AML PDX (***Supplementary Figure 10I***). Additionally, gene set enrichment analysis of the RNA sequencing data obtained from the individual AML patient samples at baseline revealed that sustained responders had a positive enrichment of gene signatures associated with translation / viral infection, and negative enrichment for gene signatures associated with cell cycle, antiviral immunity, transmembrane transport, and heme scavenging in sustained compared to poor responders to imetelstat (***Figure 6D; Supplementary Figure 9B***). Hallmark signatures revealed significant enrichment of gene sets annotated as apoptosis, interferon-alpha response, DNA repair, TP53 pathway, peroxisome, fatty acid metabolism, and reactive oxygen species pathway (***Figure 6E***).

**Figure 6:**
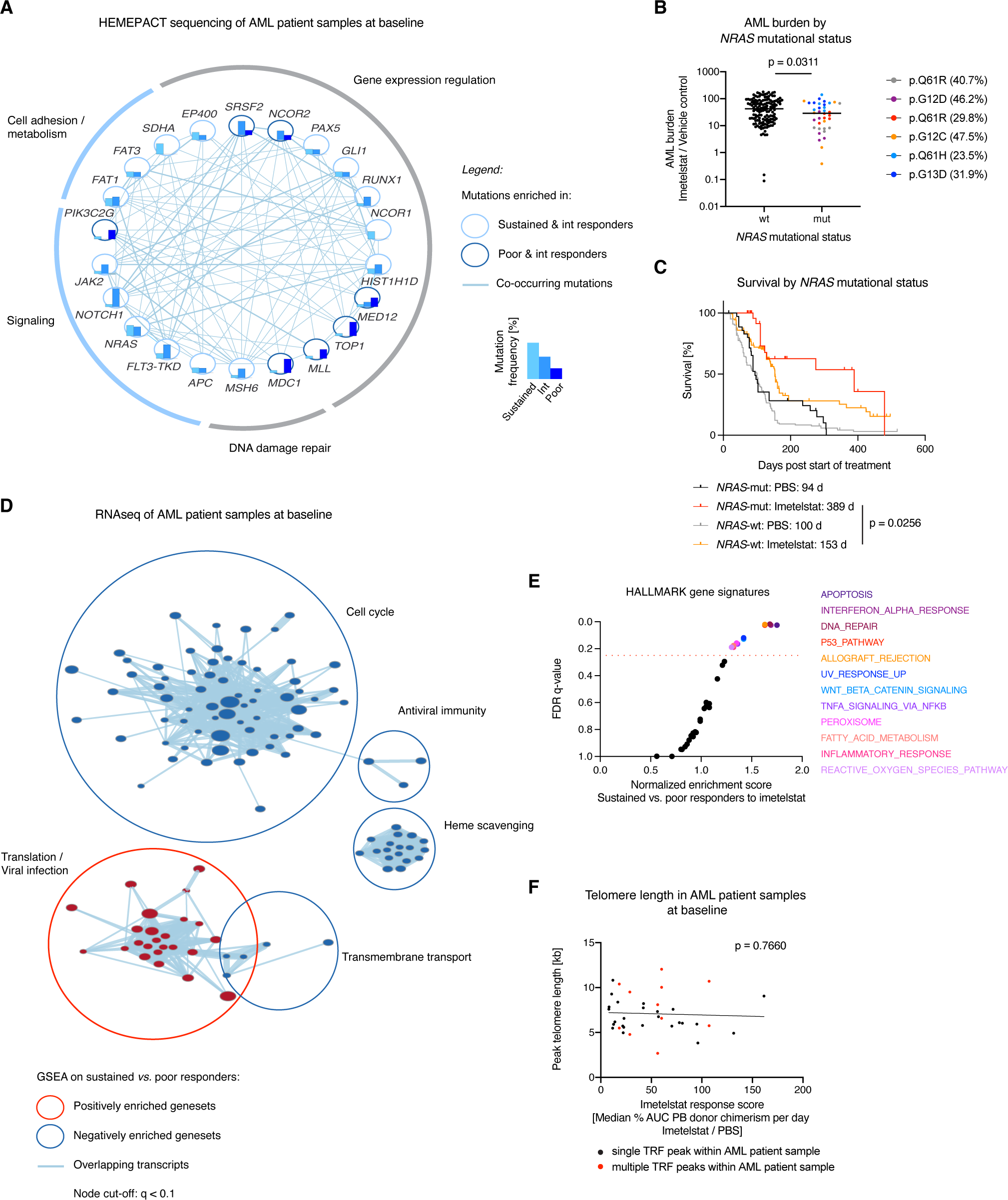
Mutant *NRAS* and oxidative stress gene expression signatures associate with sustained responses to imetelstat. Individual AML patient samples were segregated into sustained, intermediate, and poor responders to imetelstat based on peripheral blood AML burden with n = 14 (sustained), n = 8 (intermediate), n = 8 (poor). (A) Cytoscape visualization of the frequencies of genes with oncogenic mutations (based on the COSMIC database ^74^) in sustained (turquoise), intermediate (light blue), and poor (dark blue) responders to imetelstat. Connecting lines represent co-occurring mutations within the same individual AML patient sample. (B) AML burden in imetelstat-treated normalized to vehicle control-treated PDX in relation to *NRAS* mutational status: *NRAS* wild-type (wt; n = 144), mutant *NRAS* (mut; n = 36) consisting of pQ61R (40.7% variant allele frequency (VAF); n = 6), pQ61R (29.8% VAF; n = 6), p.Q61H (23.5% VAF; n = 6), p.G12C (47.5% VAF; n = 6), p.G12D (46.2% VAF; n = 6), and p.G13D (31.9% VAF; n = 6). Statistics according to two-sided Student’s t test: p = 0.0311 for *NRAS*- wt versus *NRAS*-mut. (C) Survival analysis of PBS and imetelstat-treated AML PDX divided into groups based on their *NRAS* mutation status. Median survival was 94 (PBS-treated *NRAS*-mut), 389 (imetelstat-treated *NRAS*-mut), 100 (PBS-treated *NRAS*-wt), and 153 (imetelstat-treated *NRAS*-wt) days from start of treatment. p = 0.0256 comparing imetelstat-treated *NRAS*-mut to imetelstat-treated *NRAS*-wt PDX according to Gehan-Breslow-Wilcoxon; n = 6 individual *NRAS*-mut AML patient samples, and n = 24 individual *NRAS*-wt AML patient samples; n = 6 PDX per treatment group per individual patient sample. (D) Cytoscape visualization of gene set enrichment analysis (GSEA) results on RNAseq data from individual AML patient samples at baseline comparing sustained with poor responders to imetelstat (node cut-off: q < 0.1). Red circles represent gene sets positively enriched in sustained versus poor responders to imetelstat. Blue circles represent negatively enriched gene sets in sustained *versus* poor responders to imetelstat. (E) Hallmark gene set enrichment analysis of RNAseq data comparing sustained versus poor responders to imetelstat at baseline. The red dotted line represents the cut-off considered for significant enrichment at FDR = 0.25. (F) Correlation analysis between baseline telomere length quantified by telomeric restriction fragment analysis and imetelstat response measured as AML burden in imetelstat-treated normalized to vehicle control-treated PDX.

We next examined whether baseline telomere length could predict imetelstat response. Telomere length was determined by telomere restriction fragment analyses, and peak telomere lengths varied between 2.7 and 12 kb amongst individual AML patient samples (***Supplementary Figure 10J***). Five out of the 30 AML patient samples contained multiple subclones with distinct telomere lengths (***Supplementary Figure 10J***). Overall, there was no correlation between baseline telomere length and imetelstat response (***Figure 6F***).

These data demonstrate that imetelstat is effective in a large proportion of AML PDX. Furthermore, sustained responses to imetelstat are independent of baseline telomere length, and are associated with marked improvements in survival, mutant *NRAS*, and baseline molecular signatures annotated as oxidative stress.

### Initial treatment with standard induction chemotherapy to induce oxidative stress sensitizes AML patient samples to imetelstat

The finding that responses to imetelstat are associated with baseline molecular signatures annotated as oxidative stress, and that the mechanism of action of imetelstat features ROS-mediated ferroptosis led to the hypothesis that oxidative stress induction can sensitize to imetelstat therapy.

Standard induction chemotherapy composed of cytarabine and an anthracycline is a potent inducer of ROS ^41^. To test whether oxidative stress-inducing therapy can sensitize AML cells to imetelstat treatment, we pre-treated AML cell lines with oxidative stress-inducing standard induction chemotherapy (i.e. cytarabine in combination with doxorubicin) and subsequently switched to imetelstat treatment. Standard induction chemotherapy significantly increased ROS levels in a dose-dependent manner that led to augmented cell death in AML cell lines (***Figures 7A-B***). In a pilot study using an *NRAS*-wild type AML PDX model (poor responder to imetelstat monotherapy; RBWH-44), a single dose of standard induction chemotherapy followed by a single dose of imetelstat resulted in significantly increased ROS levels in AML patient-derived cells in PDX *in vivo (****Figure 7C***). At this early timepoint, lipid peroxidation was not significantly different between the treatment groups (***Figure 7C***). However, after a complete cycle of induction chemotherapy followed by prolonged treatment with imetelstat consolidation therapy, both lipid peroxidation and ROS levels were significantly increased (***Figure 7D***).

**Figure 7:**
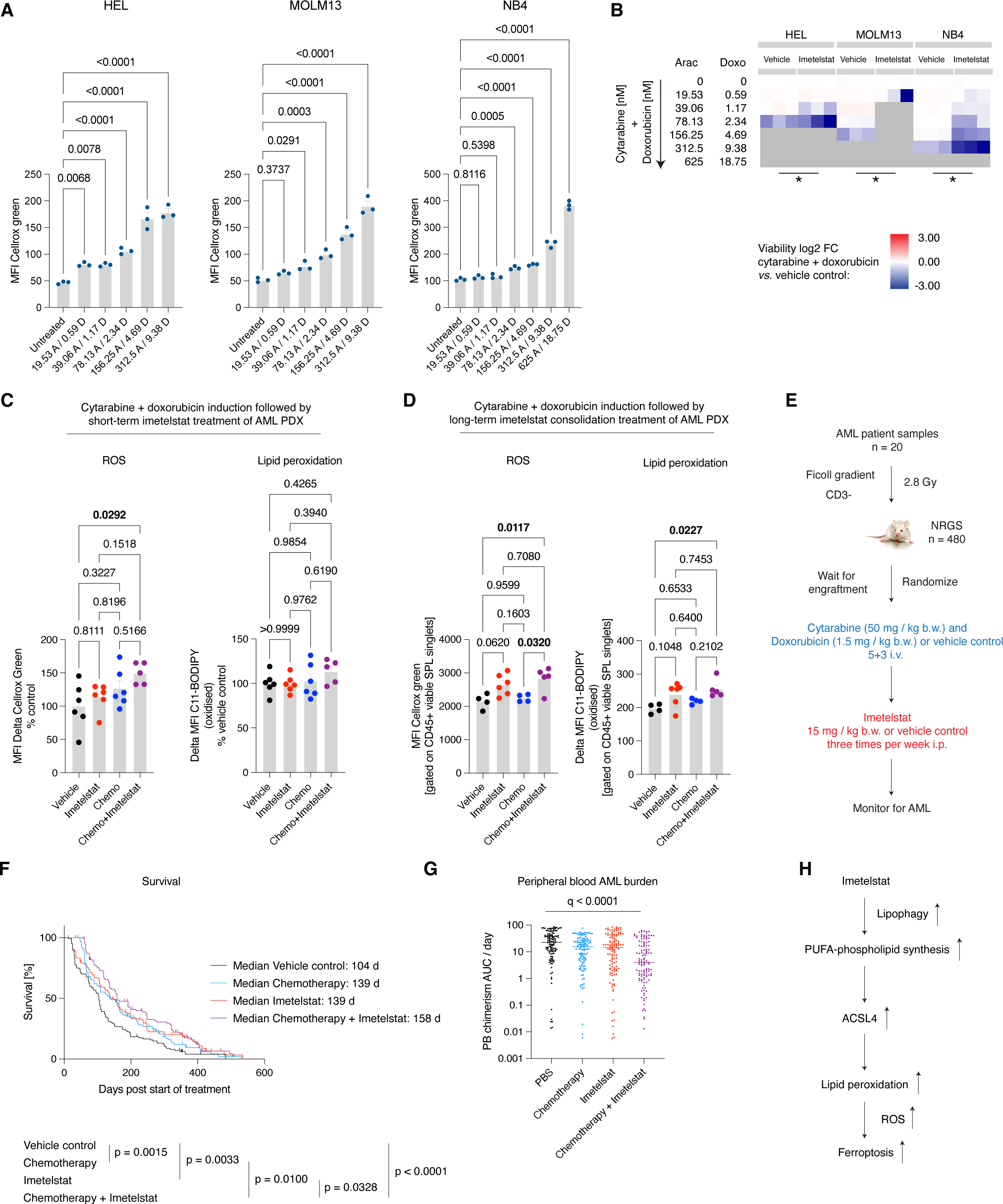
Initial treatment with standard induction chemotherapy to induce oxidative stress sensitizes AML patient samples to imetelstat. (A) Flow cytometry-based quantification of ROS levels using CellROX green in HEL, MOLM13, and NB4 cells. Data represent MFI Cellrox Green measured in chemotherapy (cytarabine plus doxorubicin) or vehicle control conditions after 3 days of treatment. N = 3 replicates. Statistics based on one-way ANOVA with multiple comparisons to untreated conditions. Adjusted p-values are displayed above each comparison. (B) Flow cytometry-based analysis of cell viability of HEL, MOLM13, and NB4 cells after switch to imetelstat-supplemented medium (4 μM). Heatmaps represent log2FC of the percentage of viable, sytox- cells in treatment conditions when compared to vehicle control. N = 3 replicates. (C-D) Lipid peroxidation and ROS measurements in AML PDX from a poor responder to imetelstat monotherapy (RBWH-44) treated with chemotherapy induction followed by imetelstat consolidation therapy compared to double vehicle or chemotherapy / imetelstat monotherapy control groups. (C) AML PDX received a single dose of cytarabine+doxorubicin chemotherapy on day 1 followed by a single dose of imetelstat on day 2 and were analyzed on day 3. (D) AML PDX received a complete cytarabine+doxorubicin cycle (5+3 regimen) followed by imetelstat consolidation and were analysed three months post-start of treatment. N=4-6 PDX per treatment group. Statistics based on One-way ANOVA with multiple comparisons as indicated. Q- values < 0.05 are highlighted in bold. (E) Experimental scheme of the sequential administration of standard chemotherapy followed by imetelstat consolidation therapy in a randomized phase II-like preclinical trial in AML patient-derived xenografts. (F) Kaplan-Meier plot showing AML-free survival for AML PDX treated with imetelstat following standard induction chemotherapy. The median survival for imetelstat following chemotherapy-treated PDX was 158 days versus 139 days (chemotherapy alone; p = 0.01), 139 days (imetelstat alone; p = 0.0328), or 104 days vehicle control (PBS) post-start of treatment with p < 0.0001 according to Gehan-Breslow-Wilcoxon. N = 120 PDX per group with 6 PDX per treatment group per individual AML patient sample, with n = 20 individual patient samples. (G) AML burden measured as peripheral blood donor chimerism (area under the curve per day) in vehicle control (PBS)-, chemotherapy-, imetelstat-, or chemotherapy followed by imetelstat therapy- treated AML PDX. Statistics according to one-way ANOVA. P.adj < 0.0001 for vehicle control vs. chemotherapy + imetelstat combination therapy group. (H) Model demonstrating the working hypothesis on imetelstat-induced ferroptosis in AML generated from this study.

Finally, as proof-of-concept *in vivo*, we sequentially administered oxidative stress-inducing standard induction chemotherapy prior to imetelstat in a diverse PDX cohort from 20 distinct AML patient samples (***Figure 7E***). Combination therapy significantly prolonged survival when compared to imetelstat monotherapy (158 days vs. 139 days; p = 0.0328), induction chemotherapy alone (158 days vs. 139 days, p = 0.0100), or vehicle control (158 days vs. 104 days, p < 0.0001; ***Figure 7F***). AML burden was significantly reduced in the combination therapy group when compared to either monotherapy or vehicle treated control groups (***Figure 7G***).

These data demonstrate that the rational sequencing of imetelstat and chemotherapy, using standard induction chemotherapy to induce oxidative stress and sensitize AML cells to imetelstat-induced lipid peroxidation and ferroptosis, results in significantly improved disease control of AML (***Figure 7H***).

## DISCUSSION

Imetelstat is a first-in class telomerase inhibitor with clinical efficacies in hematologic myeloid malignancies including essential thrombocythemia, myelofibrosis, and lower-risk myelodysplastic syndromes ^10–12^. The efficacy of imetelstat in AML and its mode of action have remained elusive to date. By developing and utilizing a comprehensive AML PDX resource and human cell lines for genomics, transcriptomics, and lipidomics approaches combined with functional genetic and pharmacological validation experiments, we demonstrate that imetelstat is a potent inducer of ferroptosis that effectively diminishes AML burden and delays relapse following chemotherapy.

Ferroptosis is a recently discovered type of non-apoptotic regulatory cell death that relies on the balance of the production of ROS during lipid peroxidation and the antioxidant system, and it is generally characterized by three hallmarks: 1) loss of peroxide repair capacity through GPX4, 2) availability of redox-active iron, and 3) oxidation of polyunsaturated fatty acid-containing phospholipids ^31, 42^. The experiments performed in this study have revealed evidence for imetelstat directly affecting the third hallmark of ferroptosis, the increased synthesis and subsequent oxidation of PUFA phospholipids. In AML PDX *in vivo*, imetelstat-induced lipid peroxidation is associated with significantly increased ACSL4 expression. In human AML cell lines, imetelstat treatment significantly increased lipid ROS levels that preceded massive cell death. Treatment with the lipid ROS scavengers ferrostatin-1 or liproxstatin-1 rescued imetelstat-induced cell death in all AML cell lines tested. In addition, pharmacological iron chelation using deferoxamine mesylate, 5- lipoxygenase inhibition using zileuton, or menadione supplementation were able to prevent imetelstat-induced cell death in a significant proportion of AML cell lines tested. In contrast to ferrostatin-1 and liproxstatin-1, higher concentrations of deferoxamine mesylate were detrimental for AML cells, suggesting that iron availability is crucial for AML cell survival at a level specific for each cell line. Iron metabolism is altered in AML at the cellular and systemic level, and elevated iron levels help to maintain the rapid growth rate of AML cells by activating ribonucleotide reductase that catalyzes DNA synthesis in an iron-dependent manner ^43^. Interestingly, imetelstat has shown efficacy in patients with pathology featuring ringed sideroblasts, a cellular morphological abnormality that is defined by iron-laden granules in mitochondria surrounding the nucleus, further supporting the role of iron-dependent cell death ^11, 12, 44^.

Our functional genetic experiments using Brunello library CRISPR/Cas9 editing have provided further evidence that imetelstat restricts leukemic progression via ferroptosis, revealing a closely related functional network of seven genes. One of the identified targets, ACSL4, has previously been identified as a key regulator of ferroptosis sensitivity through the shaping of the cellular lipid composition ^30^. We have functionally validated the most significantly enriched targets, FADS2 and ACSL4. The canonical role of FADS2 in fatty acid metabolism is the catalysis of the desaturation of linoleic and α-linolenic acid to long-chain PUFAs ^45, 46^. Lipidomics analysis has revealed increased levels of phospholipids containing triglycerides, and also increased levels of phospholipids containing PUFAs with three unsaturated bonds in imetelstat-treated AML cells in an FADS2-dependent manner. Using a fluorescent sensor, we have confirmed that imetelstat stimulates lipid peroxidation. These data demonstrate imetelstat-induced alterations in fatty acid metabolism that promote the formation of substrates for lipid peroxidation. Interestingly, in some lung cancer cell lines, FADS2 activation is associated with ferroptosis suppression ^47^. This dichotomy may be explained by the fact that in some cancer cells, FADS2 enables the desaturation of palmitate to sapienate (cis-6-C16:1) as part of an alternative desaturation pathway, thus potentially reducing the levels of monounsaturated fatty acids and ultimately PUFA-containing phospholipids as substrates for lipid peroxidation ^48^.

G-quadruplexes are recognized by and regulate the activity of many proteins involved in telomere maintenance, replication, transcription, translation, mutagenesis, and DNA recombination ^49–52^. The recognition of G-quadruplexes can be dictated by R-loops that show a close structural interplay and can modulate responses involving DNA damage induction, telomere maintenance, and alterations in gene expression regulation ^53^. Interestingly, G-quadruplex/R-loop hybrid structures were detected *in vitro* in the human *NRAS* promoter and at human telomeres ^54–57^. R-loop binders and epigenetic R-loop readers have been recently linked to altered fatty acid metabolism and ferroptosis ^58–60^. Moreover, constitutively activated RAS/MAPK signaling downstream of mutant NRAS is associated with enhanced sensitivity to ferroptosis ^42, 61, 62^, however, the activity of this pathway alone is unlikely the sole determinant of ferroptosis sensitivity ^63, 64^. Our integrative analysis of transcriptomics with functional genetics data has identified imetelstat target candidates that were recently discovered as G-quadruplex binding proteins (i.e. VIM, LMNA) ^38^. Moreover, VIM and LMNA have been characterized as proteins directly interacting with lipid droplets ^65, 66^.

Recent independent work has provided evidence for a role of lipid droplets in ferroptosis. In hepatocytes, the degradation of intracellular lipid droplets via autophagy (lipophagy) promotes RSL3-induced ferroptosis by decreasing lipid storage that subsequently induces lipid peroxidation ^38^. Our imaging flow cytometry analysis demonstrates significantly increased co-localization of markers for lipid droplets and late endosomes, proposing imetelstat-induced lipophagy as trigger for ferroptosis in AML.

Using a newly established, comprehensive AML patient-derived xenograft resource that reflects the overall genetic abnormalities found in large clinical cohorts, we demonstrated proof-of-concept for the sequential administration of standard induction chemotherapy followed by imetelstat consolidation to induce oxidative stress and sensitize AML patient samples to imetelstat treatment. This approach was able to cause significant delay or prevention of AML relapse. The efficacy of sequential therapy suggests that imetelstat may be particularly useful in preventing relapse after chemotherapy, for example, as a maintenance therapy. Recently, maintenance therapy with oral CC486 has shown a survival benefit in AML, however, there is no survival plateau and therefore, most patients still relapse and die of their disease ^67^. A significant proportion of AML patient samples tested (14 out of 30 samples) were classified as sustained responders to imetelstat monotherapy, and are characterized by genetic lesions in genes involved in cell adhesion, metabolism, and signalling, with the most striking result obtained for *NRAS. NRAS* is the fourth most commonly observed gene with driver mutations in adult AML ^3^. Moreover, AML cell clones harbouring mutant *NRAS* arise in some patients relapsing on targeted therapies, particularly FLT3 inhibition (i.e. crenolanib ^68^, gilteritinib ^69^), and BCL2 inhibition in some cases (i.e. venetoclax ^70, 71^). The demonstrated sustained responses to imetelstat in *NRAS* mutant AML patient samples raise the possibility that imetelstat may be used as salvage therapy or possibly in combination with FLT3 inhibitors or venetoclax to prolong remission and prevent relapse.

In conclusion, imetelstat is a potent inducer of ferroptosis that effectively diminishes AML burden and delays relapse following oxidative stress inducing chemotherapy. Clinical trials will address the efficacy of imetelstat in AML, and may focus on this compound as a consolidation strategy for preventing relapse, or potentially together with targeted therapies to improve outcomes in AML patients.

## ACKNOWLEDGEMENTS

We acknowledge all current and past members of the Gordon and Jessie Gilmour Leukaemia Research Laboratory, particularly, Emily Cooper, Jonathon Bolton, Gabor Pali, Emma Duce, Rebecca Austin, Therese Vu, Megan Bywater, Gerlinda Amor for technical assistance and fruitful discussions; Stanley Chun-Wei Lee from MSKCC for help with library preparation for HemePACT sequencing; Nic Waddell for bioinformatics support particularly sequencing data processing; Rebecca Fieth, Jamie Riches from Queensland University of Technology for technical support and proof reading the manuscript; Leesa Wockner and Louise Marquardt for help with statistical analyses; Grace Chojnowski, Michael Rist, Paula Hall, Tam Hong Nguyen, Lucie Leveque-ElMouttie, Yitian Ding for help with imaging flow cytometry analysis and cell sorting; the current and past members from the animal house facility, particularly Suzanne Cassidy, Jonathan Mauclair, David McNeilly; Paul Collins for technical assistance regarding next generation sequencing and routine genetic and functional analyses (STR profiling, mycoplasma testing); Clay Winterford, Andrew Masel, Ashwini Potadar for histological analyses; Richard Lock, Andrew Moore for AML patient samples; Vicki Whitehall, Wei Shi, Hong Ping for provision of cell lines and reagents; Tina M. Schnoeder, Glen Boyle, Bryan Day, Louise Purton for fruitful discussions; Geron Corp (Fei Huang, Kevin Eng) and Janssen Research & Development, LLC for the provision of imetelstat and its controls and fruitful discussions; all research nurses and patients; Janssen research funding agreement (2017-2018), NHMRC New Investigator project grant (CB; 2019-2022; GNT1157263), CSL Centenary fellowship (SWL).

## AUTHOR CONTRIBUTIONS

Conceptualization, C.B, L.J.B., L.B., F.H.H., M.M.H., G.A.K., H.A.P., O.A.W, and S.W.L; Methodology, C.B., A.H.P., T.S., J.S., V.L., B.J., G.H., and S.W.L; Software, J.S. and G.H.; Validation, C.B., A.H.P., A.P.S. and G.V.; Formal Analysis, C.B., A.H.P., G.V., T.S., J.S. and G.H.; Investigation, C.B. and S.W.L; Resources, C.B., V.L., G.A.K; Data Curation, C.B. A.H.P., A.S., G.V., T.S., J.S., G.C., V.S.K., A.P.S., Y.J., and R.H..; Writing – Original Draft, C.B. and S.W.L; Visualization, C.B. and T.S.; Supervision, C.B. and S.W.L; Project Administration, C.B. and S.W.L; Funding Acquisition, C.B. and S.W.L.

## METHODS

### Xenograft transplantation experiments

Primary AML samples were obtained from patients, after informed consent in accordance with the Declaration of Helsinki, and approved by the institutional (QIMR Berghofer) ethics committee protocol P1382 (HREC/14/QRBW/278). Ficoll density gradient was then used to recover viable mononuclear cells. Viably frozen primary AML cells were thawed and CD3-depleted with biotinylated anti-human CD3 (SK7) and biotin-binder Dynabeads (Invitrogen) and subsequently injected via the lateral tail vein into 2.8 Gy irradiated (24h before transplant) NSGS or NRGS recipients. For normal hematopoiesis studies, viable mononuclear cells were isolated from cord blood samples by ficoll density gradient, CD3-depleted as above, and subsequently enriched for CD34+ cells using the human CD34 MicroBead kit (130-046-702 MACS Miltenyi Biotec) according to the manufacturer’s instructions. 56,000 cells (donor 1) or 212,500 cells (donor 2) were injected via the lateral tail vein per irradiated NSG recipient. Please refer to supplementary methods for xenograft transplantation analysis protocols.

### Drug-Treatment Studies

All mouse experiments were approved by the institutional ethics committee protocol A11605M. NSG, NSGS or NRGS mice were treated with 15 mg/kg imetelstat (GRN163L), mismatch controls (Mismatch 1 also referred to as MM1 or GRN140833; Mismatch 2 also referred to as GGG-mismatch, MM2 or GRN142865) or vehicle control (PBS) via the intraperitoneal route for the period of time specified in the respective experiment three times per week, at least every 72h. For standard induction chemotherapy studies, cytarabine (AraC; 1 g / 10 ml isotonic water; Pfizer) and doxorubicin (Doxo; 50 mg / 25 ml saline; Pfizer) were freshly diluted with saline (sodium chloride 0.9% for irrigation; Baxter) to achieve a final concentration of 50 mg/kg body weight AraC or 1.5 mg/kg body weight Doxo in 200 ul total injection volume per recipient. Both AraC and Doxo were co-delivered intravenously (in the same syringe) on days 1 to 3, followed by intravenous injection of cytarabine alone on days 4 and 5, each in strict 24 h intervals. For chemotherapy plus imetelstat combination studies, the first imetelstat injection was administrated one day after the standard induction chemotherapy cycle was completed.

### CRISPR/Cas9 screen

The Brunello genome-wide gRNA library contains 76,441 gRNAs targeting 19,114 genes and was obtained from Addgene (Cat# 73178). Lentivirus containing the Brunello library was generated and used to transduce NB4 cells. Please refer to the supplementary methods section for detailed descriptions of the procedures.

### Lipidomics: Targeted LC/MS analysis

Targeted lipidomics was performed on a 1290 Infinity II UHPLC coupled to a 6470 QQQ mass spectrometer via AJS ESI source (Agilent, Santa Clara, USA) in positive ionization mode, using a scheduled multiple reaction monitoring (MRM) method adapted from Huynh and co-workers ^72^. The MRM transition list contained 20 lipid classes and 593 lipid species (excluding internal standards CUDA and SPLASH Lipidomix). Skyline-daily ^73^ and lipidr ^29^ software were used for data analysis. Please refer to the supplementary methods section for detailed descriptions of the procedures.

### Data availability

All Skyline and mass spec raw data of targeted lipidomics experiments have been deposited on Panorama Public (https://panoramaweb.org/ImetelstatLipidomics.url) [ and Password: tEimcwMc].

RNAseq datasets have been reposited at GEO:

GSE176522 - https://www.ncbi.nlm.nih.gov/geo/query/acc.cgi?acc=GSE176524 [Reviewer access token: urexcomqjtwfdur]

GSE176523 - https://www.ncbi.nlm.nih.gov/geo/query/acc.cgi?acc=GSE176523 [Reviewer access token: ijmrmmwohxkbnmt]

