## Supplementary figures and images for "Imetelstat-Mediated Alterations in Fatty Acid Metabolism To Induce Ferroptosis As Therapeutic Strategy for Acute Myeloid Leukemia"

### Supplemental Figure 1

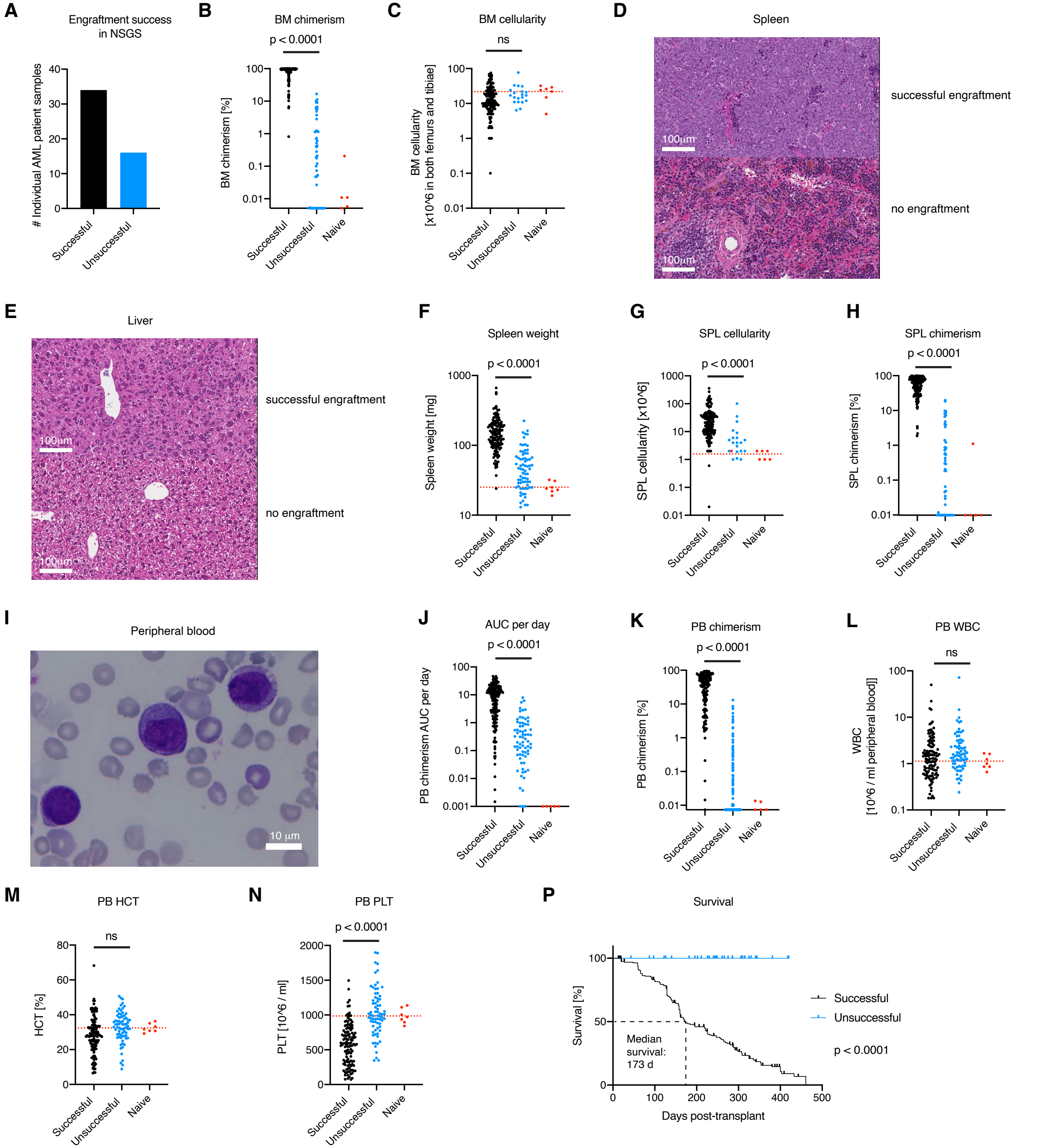


**Supplementary Figure 1**

### Supplemental Figure 2

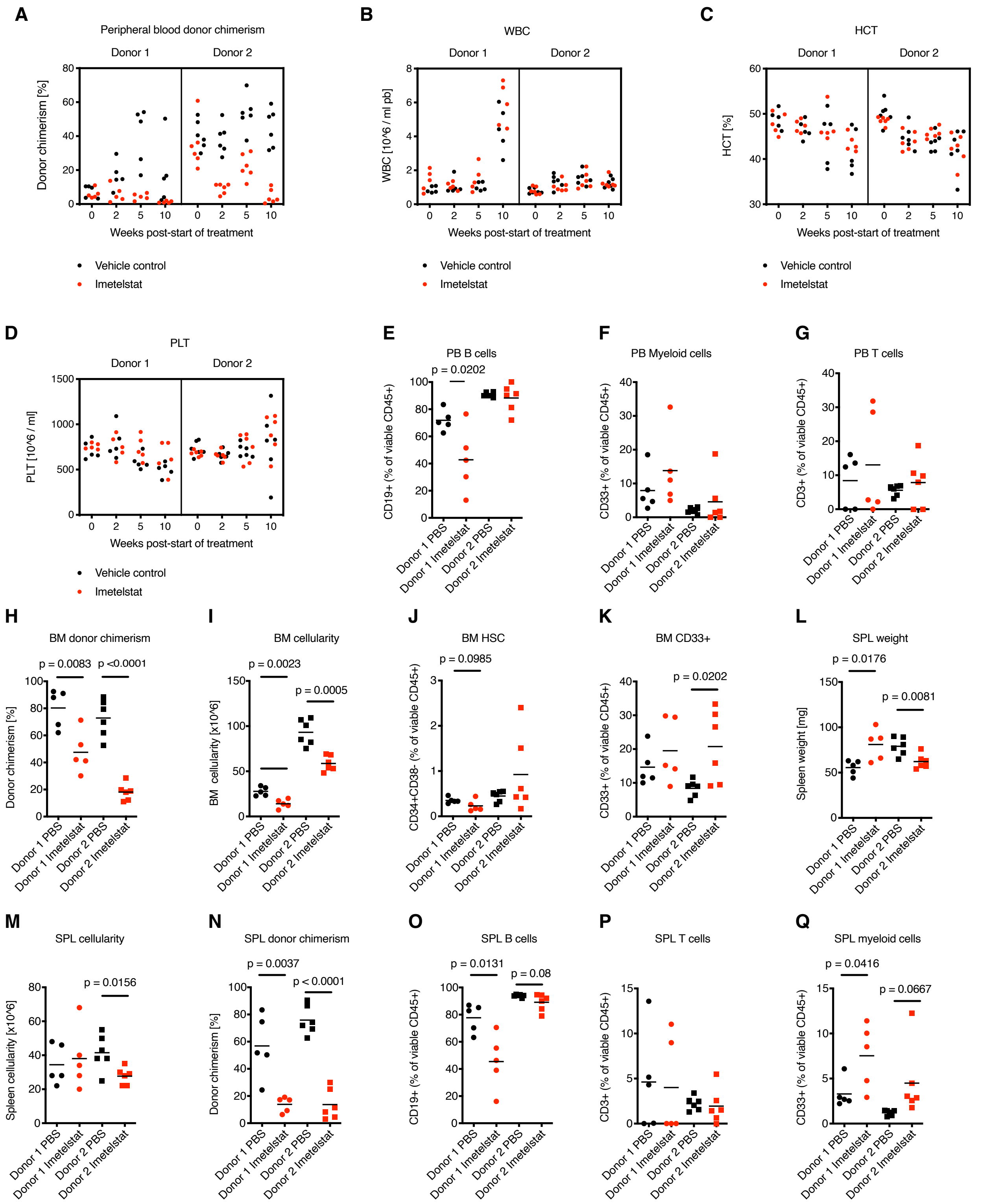


**Supplementary Figure 2**

### Supplemental Figure 3

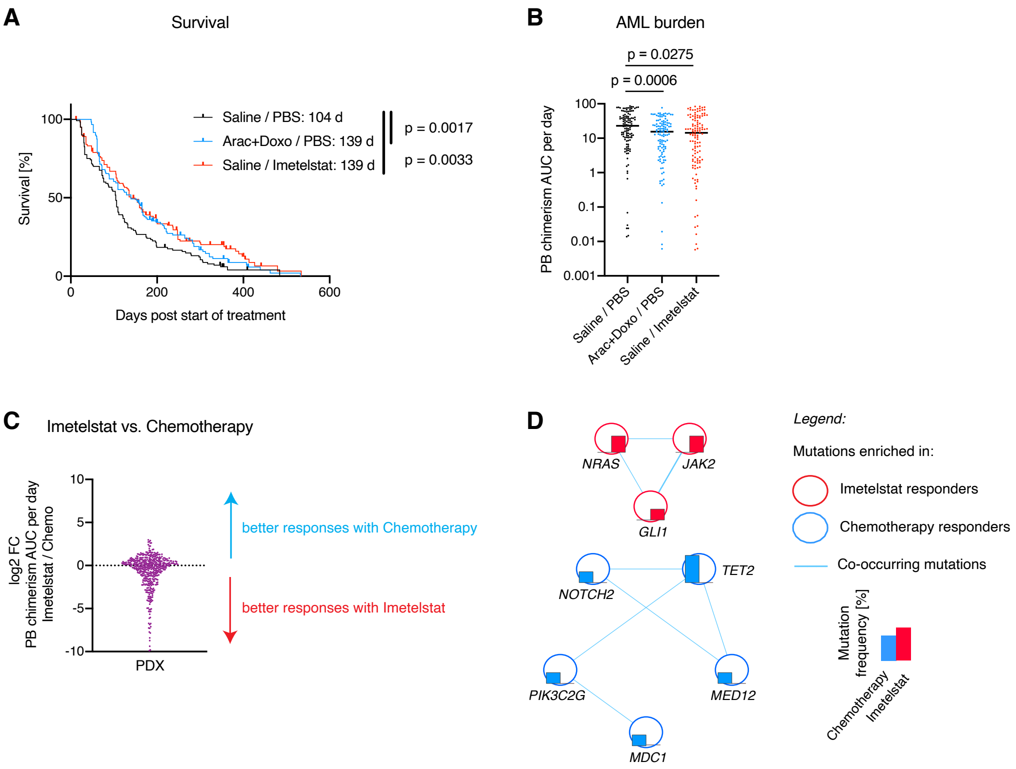


**Supplementary Figure 3**

### Supplemental Figure 4

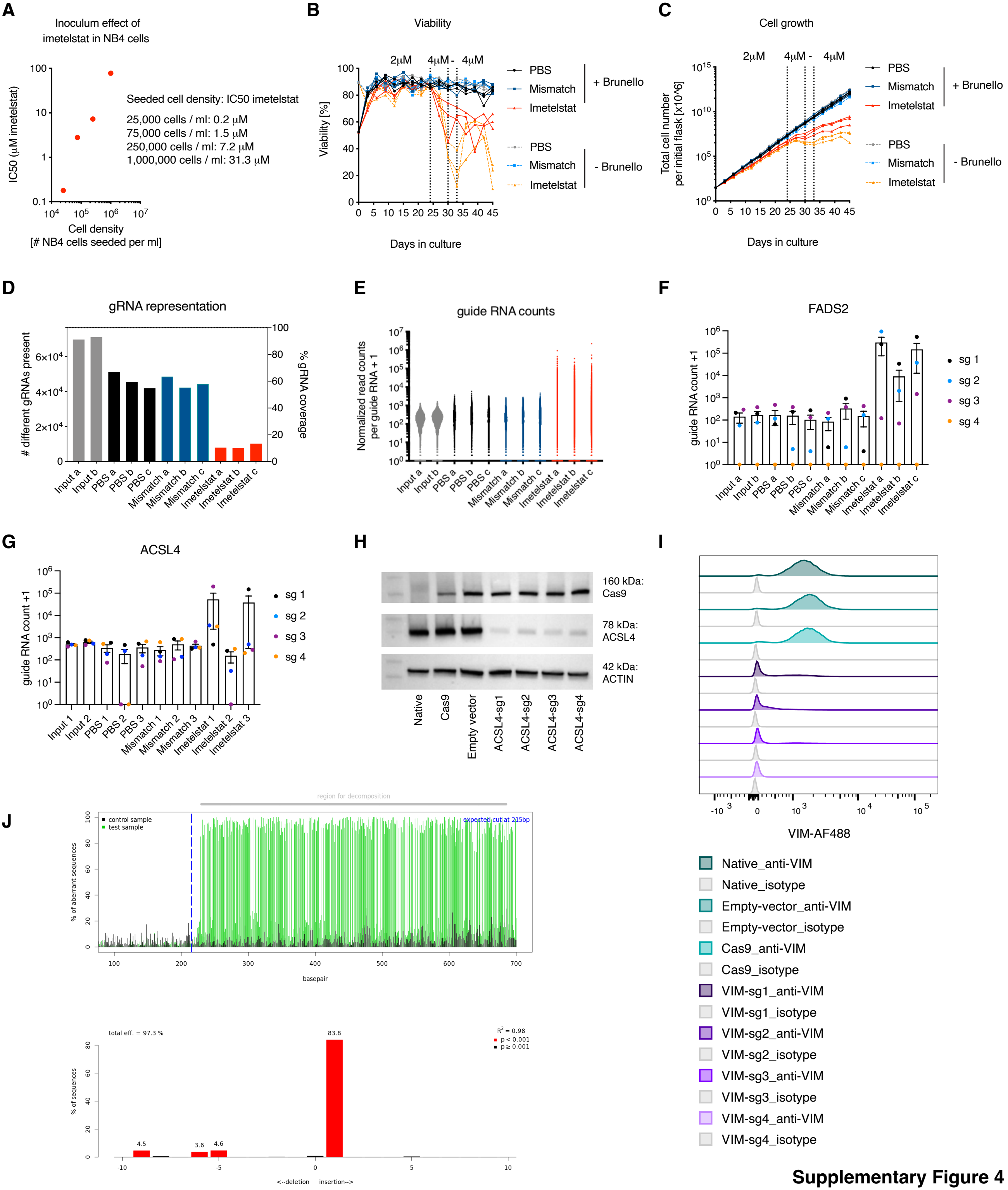


**Supplementary Figure 4**

### Supplemental Figure 5

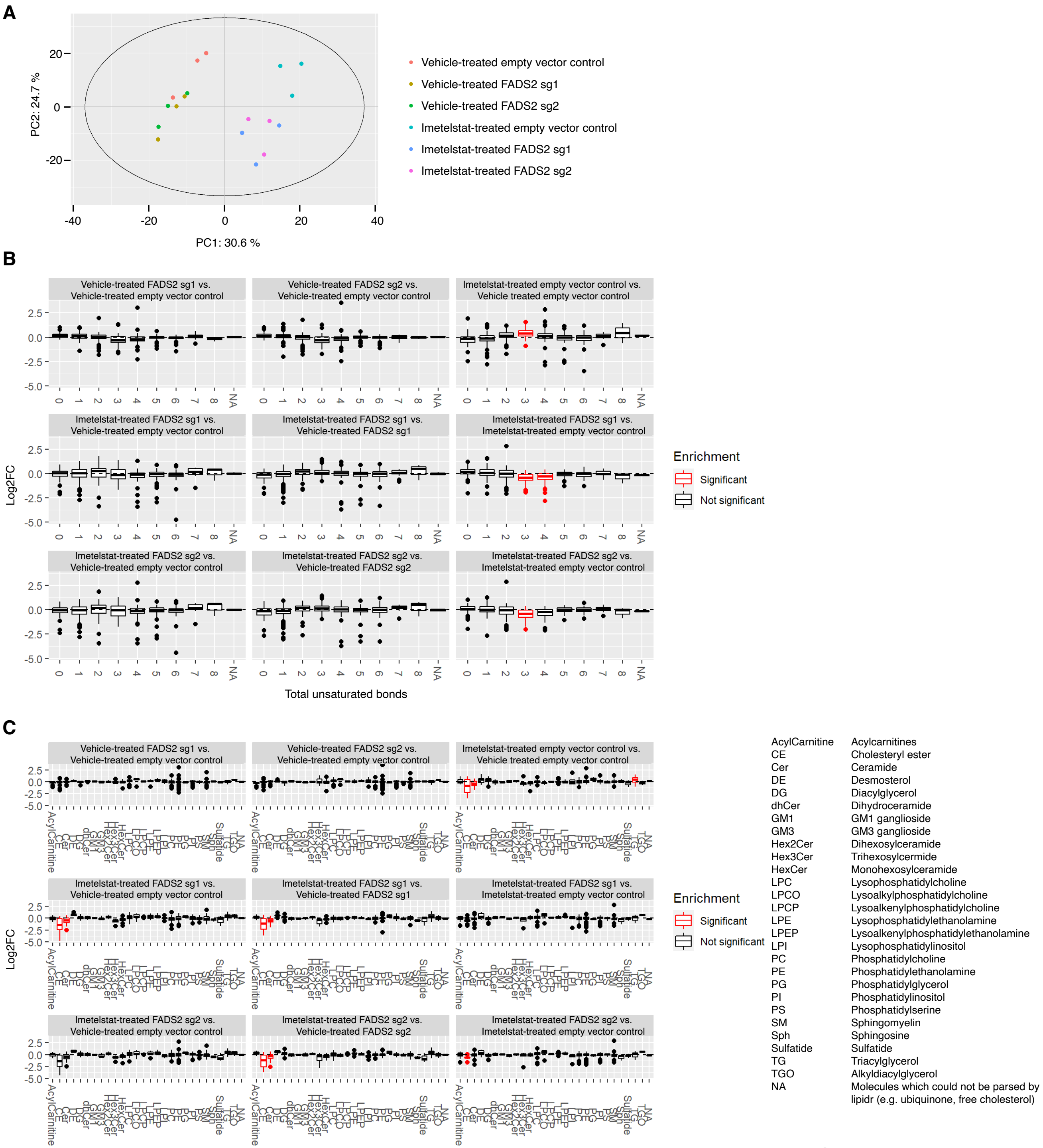


**Supplementary Figure 5**

### Supplemental Figure 6

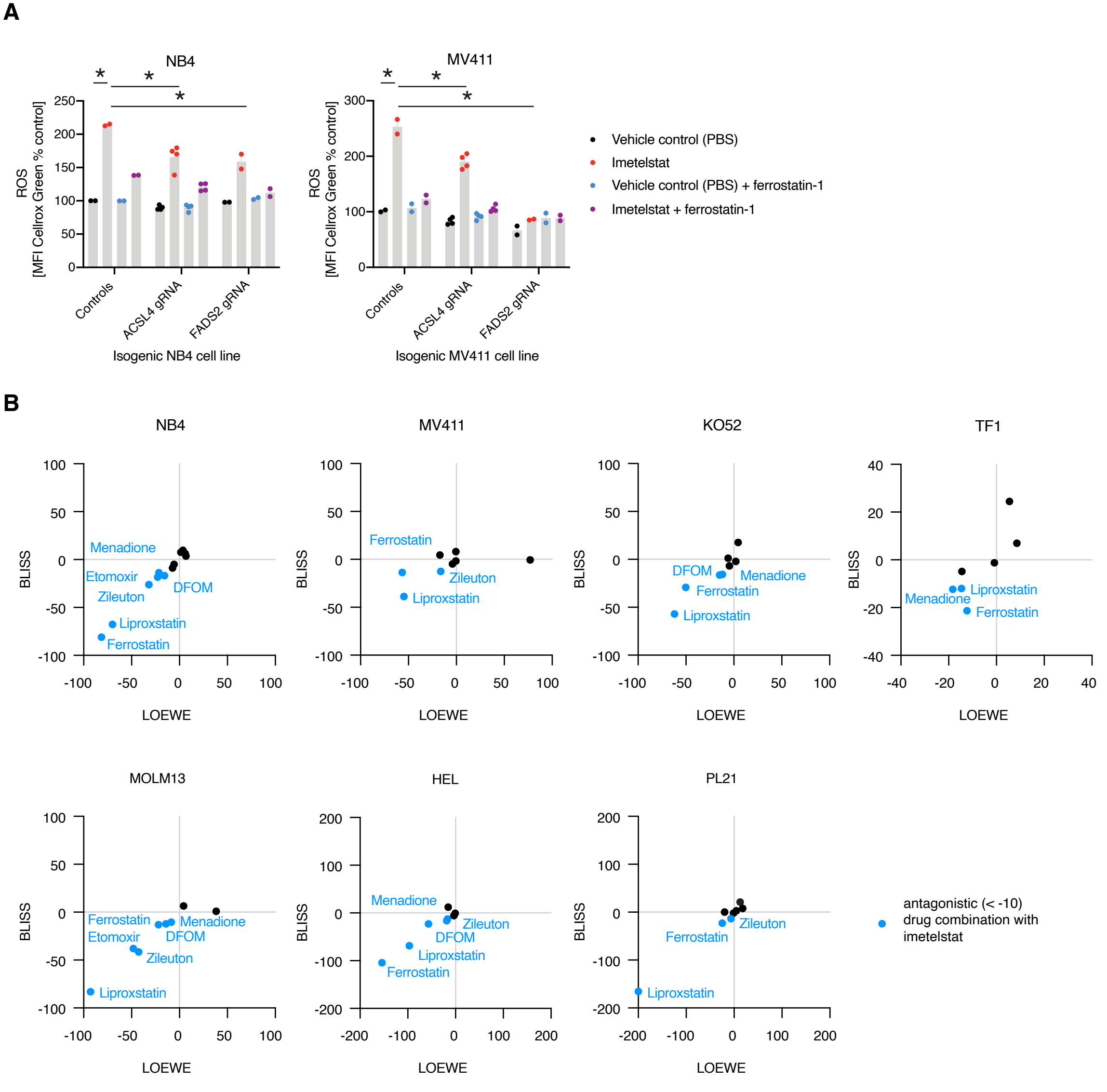


**Supplementary Figure 6**

### Supplemental Figure 7

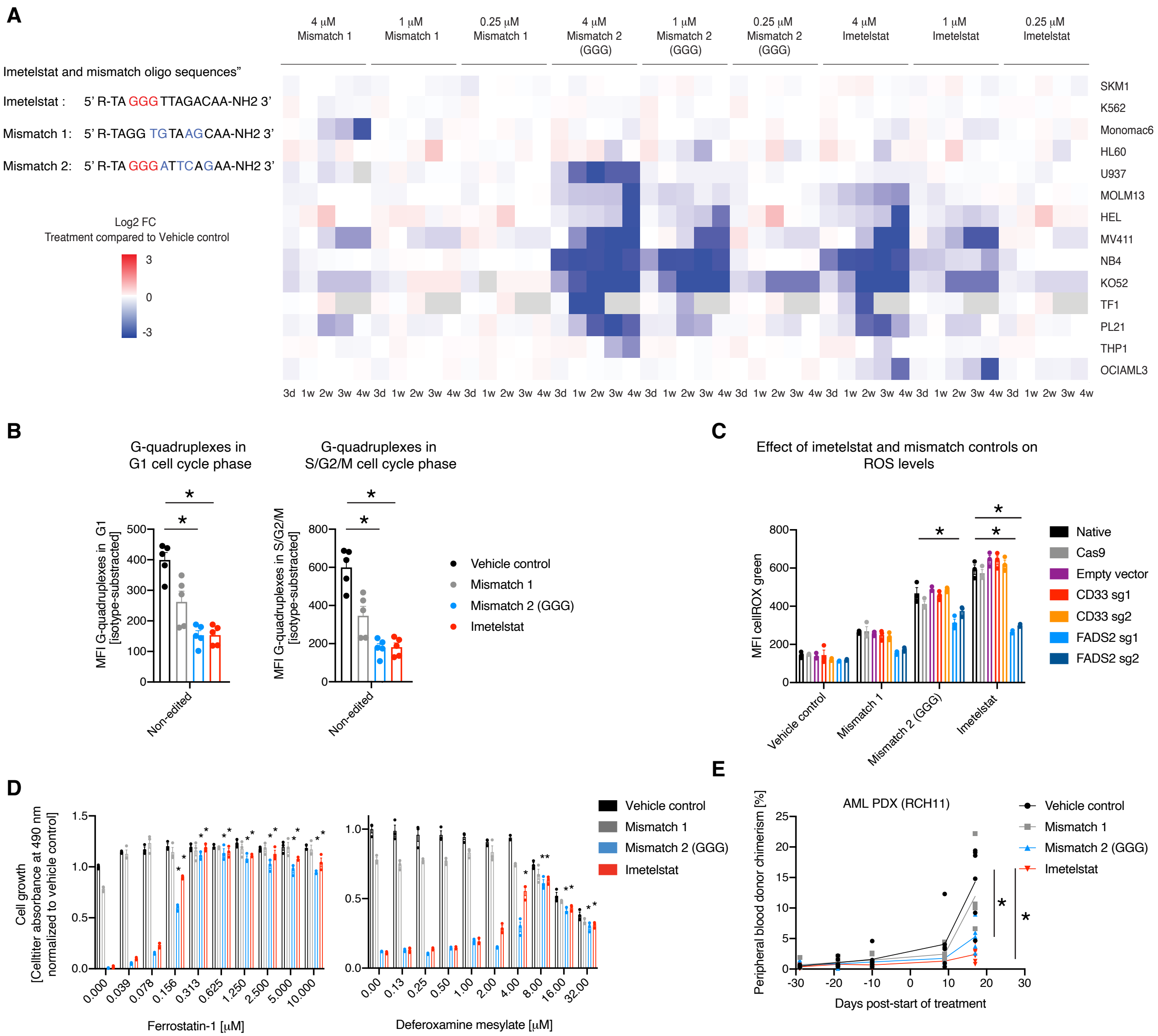


**Supplementary Figure 7**

### Supplemental Figure 8

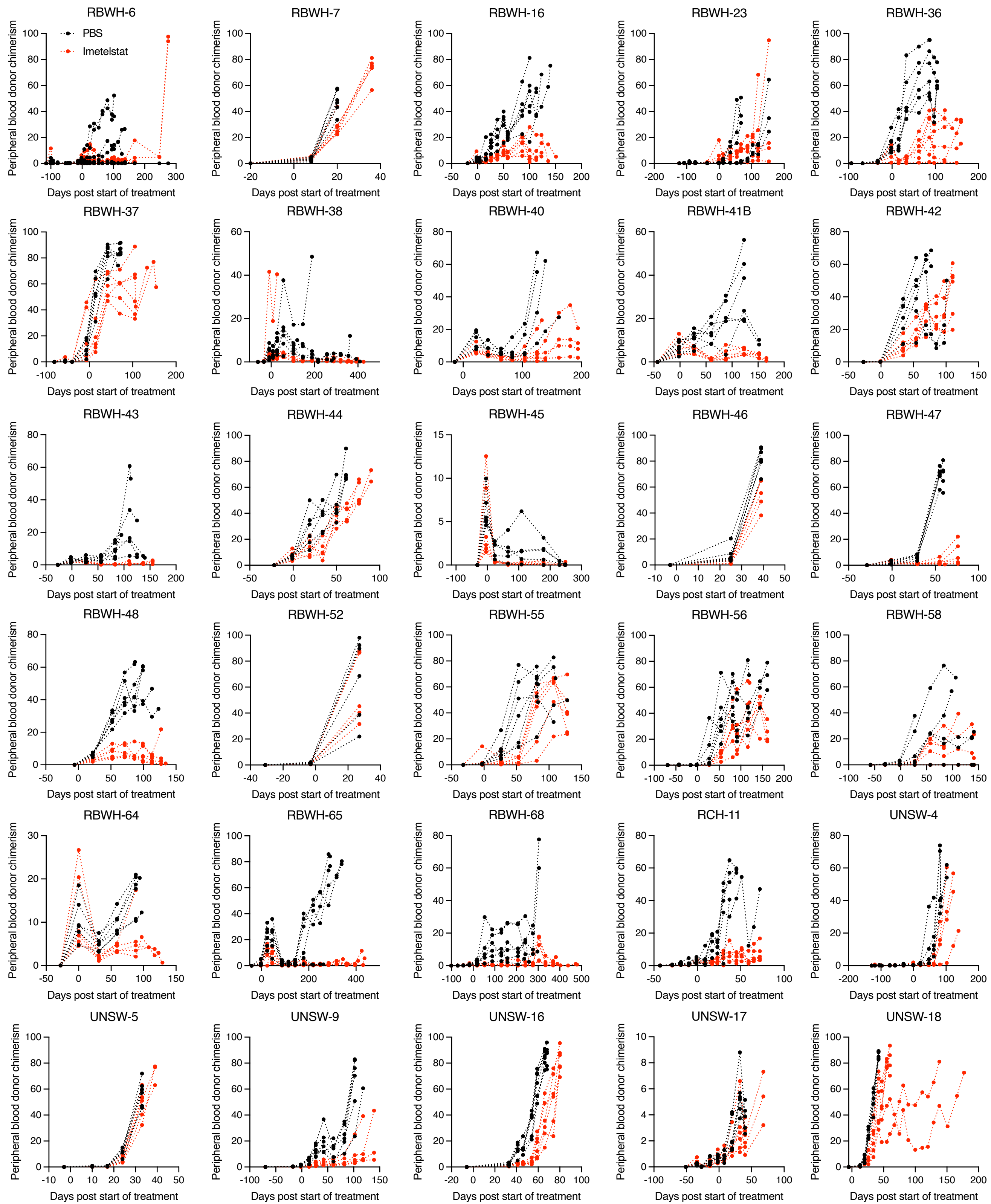

### Supplemental Figure 9

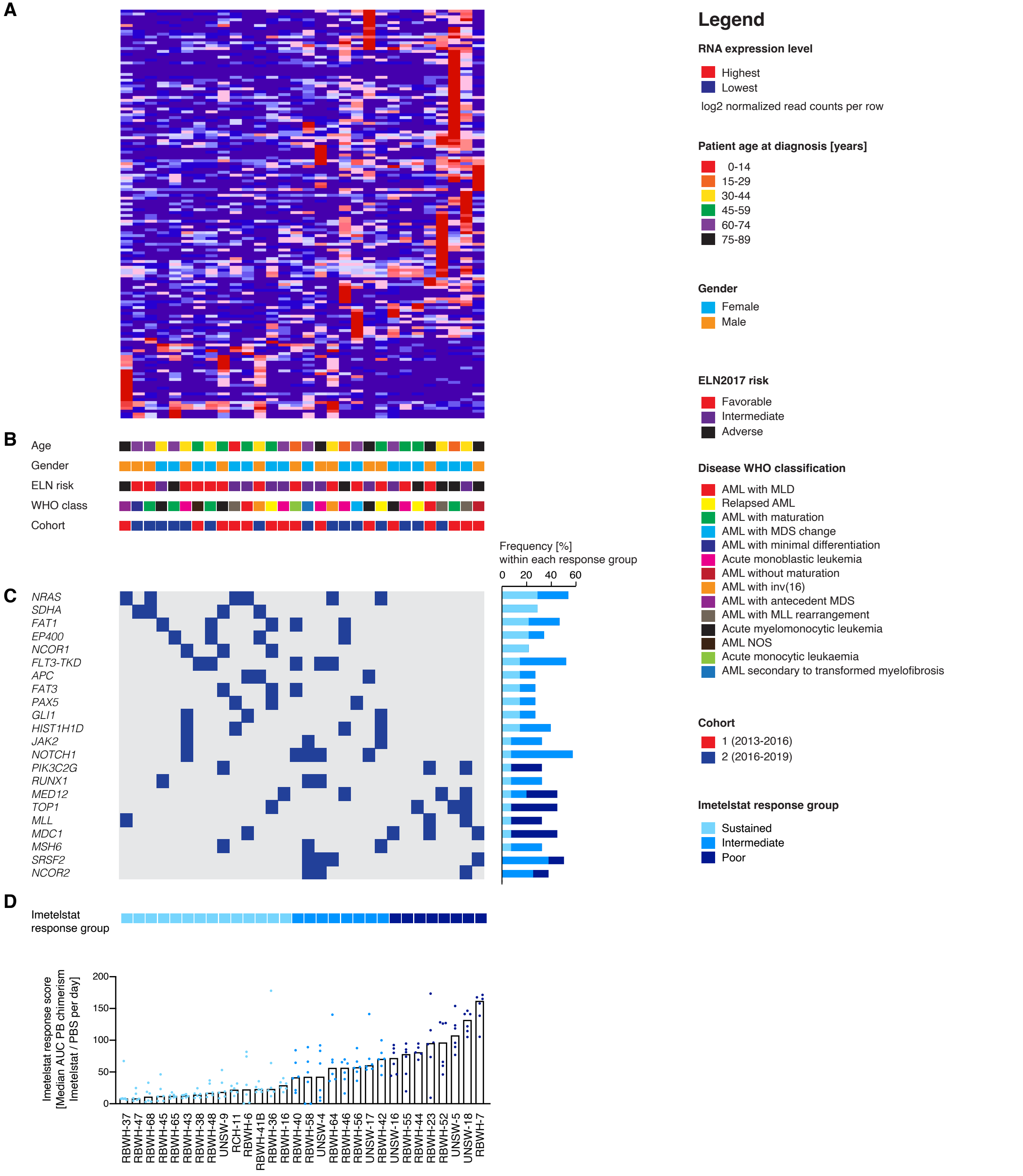


**Supplementary Figure 9**

### Supplemental Figure 10

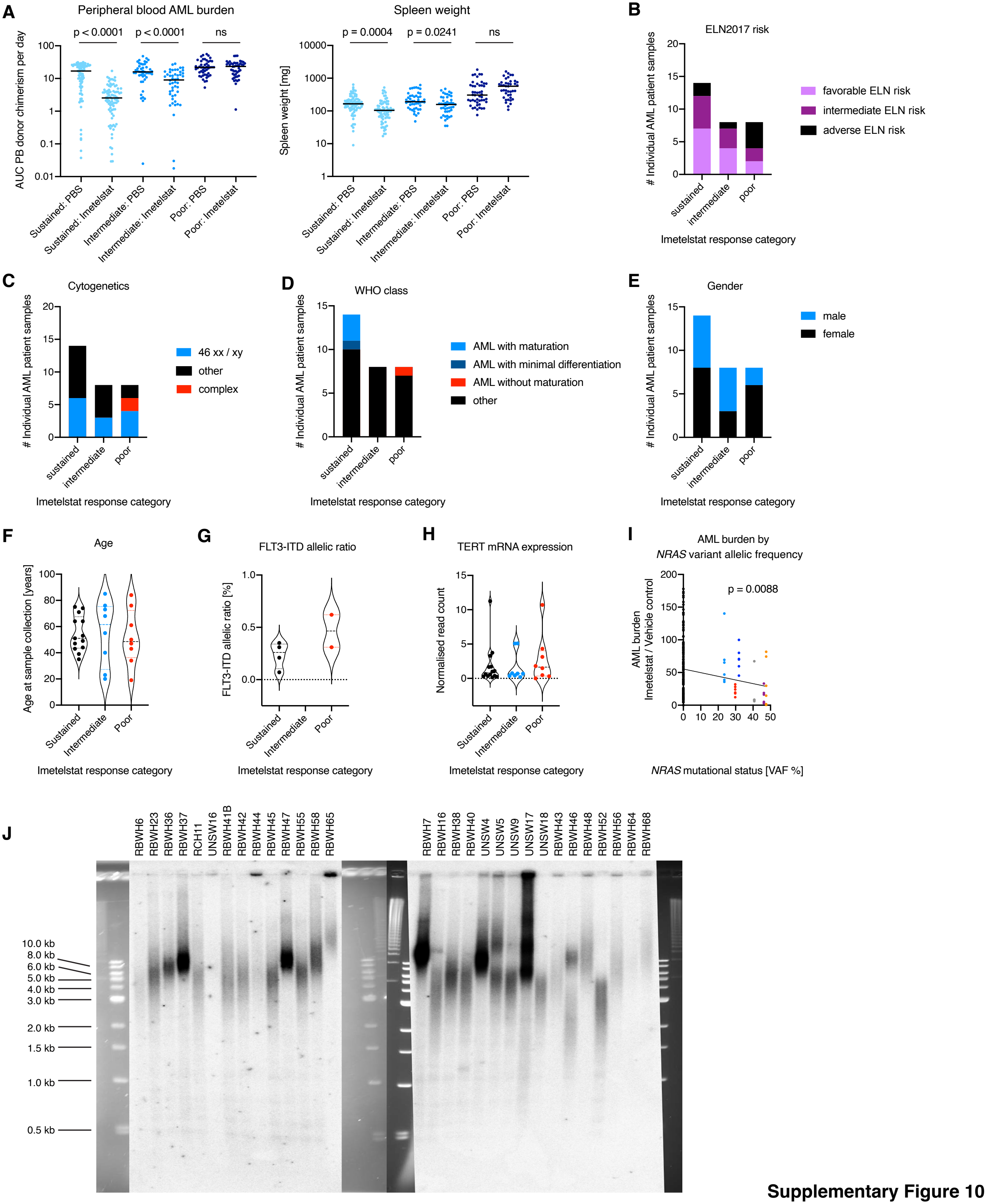


**Supplementary Figure 10**
