## Supplemental Figure Legends and Materials & Methods for "Imetelstat-Mediated Alterations in Fatty Acid Metabolism To Induce Ferroptosis As Therapeutic Strategy for Acute Myeloid Leukemia"

### SUPPLEMENTARY FIGURE LEGENDS AND MATERIALS & METHODS

**Figure S1 (related to Figure 1): Generation of a Comprehensive and Representative AML PDX Resource**

(A) The number of AML patient samples that either successfully (black) or unsuccessfully (blue) generated AML PDX using NSGS[1] recipients. (B-C) Bone marrow donor chimerism (hCD45+ % from hCD45+ and mCd45.1-; B) and cellularity (x10^6 harvested from both femurs and tibiae; C). (D-E) Histologic analysis of spleen (D) and liver (E) morphology in AML PDX. (F-H) Spleen weight (F), spleen cellularity (G), and splenic donor chimerism (H). I-N) Peripheral blood blast morphology analysis at takedown using Wright-Giemsa staining (I), donor chimerism area under the curve (AUC) per day (J), donor chimerism at takedown (K), and white blood cell counts at takedown (L). Hematocrit percentage (M) and platelets (x10^6/mL; N) from peripheral blood at takedown. P) AML-related survival analysis of successfully *versus* unsuccessfully generated AML PDX. Median survival was 173 days post-transplant in successfully generated AML PDX *versus* not reached for unsuccessfully generated AML PDX (follow-up > 365 days). $p < 0.0001$, Gehan-Breslow-Wilcoxon test; n = 196 (successful), n = 73 (unsuccessful) AML PDX. N = 7 naïve NSGS (40 weeks old) were used as controls. Significant p values < 0.05 are displayed in each panel.

**Figure S2 (related to Figure 2): The effect of imetelstat treatment on normal hematopoiesis**

Humanized *in vivo* models of hematopoiesis were generated by transplanting viable CD34+ mononuclear cells isolated from cord blood samples from two independent donors into twelve NSG recipients each (donor 1: 56,000 cells per NSG; donor 2: 212,500 cells per NSG). Recipients were treated for 10 weeks with imetelstat (15 mg/kg body weight) three times per week starting one month after transplantation or vehicle control (n = 6 PDX per treatment group per donor). (A-D) Peripheral blood time course analysis of donor chimerism (A), white blood cell counts (WBC; B), hematocrit (HCT, C), and platelet levels (PLT; D). (E-G) Flow cytometry analysis of peripheral blood for B cell surface marker expression (CD19; E), myeloid surface marker expression (CD33; F), and T cell surface marker expression (CD3; G) at 10 weeks post-start of treatment. (H-K) Bone marrow analysis of cord blood recipients at 10 weeks post-start of treatment: donor chimerism (H), cellularity (I), hematopoietic stem cell population percentage (CD34+CD38- %; J), and myeloid population percentage (CD33+; K). (L-Q) Analysis of cord blood recipient's spleens at 10 weeks post-start of treatment: spleen weight (L), cellularity (M), donor chimerism (N), B cell population (CD19+ %; O), T cell population (CD3+ %; P), and myeloid cell population (CD33+ %; Q). Statistics based on Student's t test. Significant p values < 0.05 are displayed in each panel.

**Figure S3 (related to Figures 1 and 7): Comparison of imetelstat to standard chemotherapy efficacy in AML PDX**

Comparative analysis of PDX responses to imetelstat versus standard induction chemotherapy, i.e. cytarabine (Arac; 50 mg/kg) and doxorubicin (Doxo; 1.5 mg/kg) 5+3 dose regimen (single cycle), in NRGS. (A) Median survival was 104 (vehicle control (Saline / PBS) – treated PDX; black) versus 139 (chemotherapy-treated PDX; Arac+Doxo; blue; p = 0.0017) and 139 (imetelstat-treated PDX; Saline / Imetelstat; red; p = 0.0033). Statistics based on Gehan-Breslow-Wilcoxon. N = 6 PDX per individual AML patient sample (i.e. 20) per treatment group (i.e. 120 PDX per treatment group). (B) AML burden quantified as peripheral blood donor chimerism per day. Statistics were based on ordinary One-way ANOVA with multiple comparisons. N = 6 PDX per individual AML patient sample (i.e. 20) per treatment group (i.e. 120 PDX per treatment group). Significant adjusted p values < 0.05 are displayed in each panel. (C) Imetelstat versus chemotherapy response calculation as log2FC of peripheral blood chimerism area under the curve per day in imetelstat-treated PDX *versus* chemotherapy - treated PDX. The red arrow indicates samples defined as preferential imetelstat responders, and conversely, the blue arrow indicates PDX defined as preferential chemotherapy responders. (D) Cytoscape visualization of genes with mutations identified exclusively in preferential imetelstat responders (red) or chemotherapy responders (blue) at baseline.

**Figure S4 (related to Figure 3): Identification of key mediators of imetelstat efficacy using a genome-wide CRISPR/Cas9 screen**

(A) Identification of an imetelstat inoculum effect by quantification of IC50 values in NB4 cultures seeded at different densities: IC50 = 0.2 μM (25,000 cells per ml); IC50 = 1.5 μM (75,000 cells per ml); IC50 = 7.2 μM (250,000 cells / ml); IC50 = 31.3 μM (1,000,000 cells / ml). (B-C) Viability (B) and cell growth (C) of Brunello-library transduced NB4 cells cultured in the presence of vehicle control (PBS), mismatch control (MM1), or imetelstat compared to non-transduced NB4 cells over a time course of 45 days. The respective concentrations are indicated in the panel. N = 3 replicates per genotype and treatment condition. (D-E) Next generation sequencing of DNA isolated from Brunello library-transduced NB4 cells harvested at day 45 in culture treated with vehicle control (PBS; black bars), mismatch control (MM1; blue bars), or imetelstat (red bars), and compared to DNA isolated from Brunello library-transduced NB4 cells before treatment as input control (grey bars): (D) The number of different guide RNAs present (left y-axis) and percentage of guide RNA coverage (right y-axis); (E) the read counts obtained from each guide RNA. (F-G) Presentation of the read counts obtained from *FADS2* (F) or *ACSL4* (G) targeting guide RNAs in the input, vehicle (PBS), mismatch (MM1) or imetelstat-treated Brunello-transduced NB4 cultures. Each gene was targeted by 4 distinct guide RNAs (sg1-4). (H-J) Confirmation of efficient CRISPR/Cas9 - generated knockdowns in human AML cell lines by ACSL4 western blotting (H), flow cytometric analysis of intracellular VIM expression (I), and *FADS2* gene editing by TIDE analysis (J).

**Figure S5 (related to Figure 4): Lipidomics analysis of imetelstat-treated *FADS2*-edited or non-edited NB4 cells**

Targeted lipidomics analysis on 593 lipid species and their desaturation levels. (A) PCA plot on normalized peak areas of lipid species in *FADS2*-edited (i.e. FADS2 sg1, FADS2 sg2) or non-edited (i.e. empty vector control) NB4 cultures supplemented for 24h with either 4 μM imetelstat or vehicle control. (B) Differential enrichment of unsaturated bonds in all measured lipid species. (C) Differential enrichment of lipid species according to lipid classes.

Red box plots denote significantly different comparisons with $p < 0.005$. N = 3 independent replicates per condition from distinct cell passages.

**Figure S6 (related to Figure 4): Lipid ROS are essential for imetelstat's mechanism of action in AML**

(A) Quantification of reactive oxygen species (ROS) levels using CellROX green in in *ACSL4*-edited (i.e. ACSL4-sg1, ACSL4-sg2, ACSL4-sg3, ACSL4-sg4) or *FADS2*-edited (i.e. FADS2-sg1, FADS2-sg2) or non-edited (i.e. Cas9, empty vector) NB4 or MV411 cells treated with imetelstat (4 μM) or vehicle control for 24h. Cellrox Green incubation medium was supplemented with vehicle or 500 nM ferrostatin-1 to scavenge lipid ROS during detection. Median fluorescent intensities (MFI) were defined statistically significantly different in each condition based on two-sided Student's t test with $p < 0.001$, corrected for multiple testing according to Bonferroni; n = 3 independent replicates. Asterisks (*) denote statistically significantly different comparisons to the respective vehicle control treatment condition. (B) Celltiter-based cell growth analysis of imetelstat-sensitive cell lines NB4, MV411, KO52, TF1, MOLM13, HEL or PL21 that were supplemented with different concentrations of imetelstat combined with various pharmacological modulators of ferroptosis (i.e. ferrostatin-1, liproxstatin-1, DFOM, zileuton, menadione, etomoxir, RSL3, erastin). Synergism was estimated using the Synergyfinder 2.0 algorithm (http://synergyfinder.fimm.fi), BLISS and LOEWE synergy scores are plotted for each drug combination and cell line. Ferroptosis modulators highlighted in blue represent predicted antagonistic combinations with imetelstat with LOEWE and BLISS scores < 10.

**Figure S7 (Related to Figure 5): Efficacy of imetelstat and its triple G-containing mismatch control in AML**

(A) Oligonucleotide sequences of imetelstat, mismatch 1, and mismatch 2 (GGG). Celltiter analysis of 14 human hematopoietic cell lines treated with different concentrations (0.25 μM, 1 μM, 4 μM) of mismatch 1, mismatch 2, or imetelstat at multiple timepoints during a 4-week period. N = 3 technical replicates per condition from 3 independent experiments. (B) Flow cytometric analysis of G-quadruplex structures in non-edited (i.e. native, Cas9, empty vector, CD33 sg1, CD33 sg2) NB4 cells that were treated with vehicle control, mismatch 1, mismatch 2 (GGG), or imetelstat, and gated on G1 cell cycle phase (I), or S/G2/M cell cycle phases. Each dot represents the mean of 3 technical replicates. Statistics were carried out using two-sided Student's t test corrected for multiple testing using Bonferroni. Asterisks (*) denote statistically significant differences with $p < 0.05$. (C) CellROX green measurement of unedited (i.e. native, Cas9, empty vector, CD33 sg1, CD33 sg2) or FADS2-edited (i.e. FADS2 sg1, FADS2 sg2) NB4 cells that were treated with vehicle control, mismatch 1, mismatch 2 (GGG), or imetelstat. N = 3 replicates per group. Asterisks (*) denote statistically different comparisons with $p < 0.05$ corrected for multiple testing by Bonferroni. (D) Celltiter analysis on NB4 cells treated with vehicle control, mismatch 1, mismatch 2 (GGG), or imetelstat, and in combination with ferrostatin-1 (left graph) or DFOM (right graph). N = 3 replicates per condition. Statistics according to two-sided Student's t test comparing ferrostatin-1 or deferoxamine mesylate supplemented mismatch 2 or imetelstat - treated conditions to those treated with mismatch 2 or imetelstat alone. Asterisks (*) denote statistically significant differences with $p < 0.05$ corrected for multiple testing using Bonferroni. (E) Peripheral blood donor chimerism timecourse analysis of an *NRAS/KRAS* mutant AML PDX model (RCH11) that was treated with vehicle control, mismatch 1, mismatch 2 (GGG), or imetelstat. N=6 replicates per treatment group.

**Figure S8 (Related to Figures 2 and 6): Individual AML PDX responses to imetelstat**

Peripheral blood AML donor chimerism in the thirty individual PDX models. N = 6 NSGS per treatment group per individual AML patient sample. Kinetics are presented from each individual replicate.

**Figure S9 (related to Figures 1, 2 and 6): The molecular landscapes underlying imetelstat responses in AML PDX**

(A) Unsupervised hierarchical clustering analysis on the expression of differentially expressed transcripts in sustained versus poor responders to imetelstat with a cut-off of adjusted p-value < 0.25 among 30 individual AMLs from our PDX repository. (B) Key clinical characteristics of patients from whom AML samples were derived including age at diagnosis, gender, ELN2017 prognostic risk group, and WHO class of disease. (C) OncoPrint of differentially detected mutations in AMLs from sutained versus poor responders to imetelstat at baseline by targeted next generation sequencing of 585 genes associated with hematologic malignancies (the MSKCC HemePACT assay)[2]. (D) Imetelstat response classification: AML burden quantified as area under the curve of peripheral blood donor chimerism per day of imetelstat / vehicle (PBS)-treated AML PDX generated from each individual AML patient sample. N = 30 AML patient samples with n = 6 PDX per treatment group.

**Figure S10 (related to Figure 5): Imetelstat response classification in AML PDX**

(A-B) Imetelstat response classification: AML burden quantified as peripheral blood donor chimerism per day and spleen weights from imetelstat vs. vehicle control-treated PDX by imetelstat response group. Statistics based on two-sided Student's t test on log-transformed data. N = 84 (sustained responders per treatment group each); n = 48 (intermediate or poor responders per treatment group each). Solid lines represent the median from each group. Significant p values < 0.05 are displayed in each panel. (B-H) Representation of clinical and molecular parameters in AML patient samples either characterized as sustained, intermediate, or poor responders to imetelstat: ELN 2017 risk (B), cytogenetics (C), WHO disease classification (D), gender (E), AML patient age at sampling (F), FLT3-ITD allelic ratio (G), and (H) TERT mRNA expression levels at baseline from RNAseq analysis. Results were not statistically different (i.e. according to Fisher's exact test for B-E; one-way ANOVA for F-H). N = 14 sustained, n = 8 intermediate, n = 8 poor responders to imetelstat. (I) AML burden in imetelstat-treated normalized to vehicle control-treated PDX in relation to *NRAS* mutational status: *NRAS* wild-type (wt; n = 144), mutant *NRAS* (mut; n = 36) consisting of pQ61R (40.7% variant allele frequency (VAF); n = 6), pQ61R (29.8% VAF; n = 6), p.Q61H (23.5% VAF; n = 6), p.G12C (47.5% VAF; n = 6), p.G12D (46.2% VAF; n = 6), and p.G13D (31.9% VAF; n = 6). Simple linear regression indicates the slope being significantly different from 0 with p = 0.0088. (J) Telomeric restriction fragment analysis of genomic DNA isolated from viable AML patient cells at baseline from all 30 individual AML patient samples included in the preclinical trial of imetelstat in AML PDX.

### MATERIALS & METHODS

#### Mouse Models

NSG (i.e., NOD.Cg-Prkdcscid Il2rgtm1Wjz /SzJ), NSGS (i.e., NOD.Cg-Prkdcscid Il2rgtm1Wjl Tg[CMV-IL3,CSF2,KITLG]1Eav/MloySzJ), and NRGS mice (i.e., NOD.Cg-Rag1tm1Mom Il2rgtm1Wjl Tg[CMV-IL3,CSF2,KITLG]1Eav/J) were imported from Jackson Laboratories. All mice were kept pathogen-free in the animal facility of QIMR Berghofer Medical Research Institute. Mice received autoclaved and Baytril-treated (100 mg/L; Provet) water until 1 to 7 days before irradiation, and after that autoclaved and Septrin- treated (12 ml/L cherry-flavored paediatric suspension, i.e. 96 mg/L trimethoprim and 480 mg/L sulphamethoxazole; Arrow Pharmaceuticals) water. All mouse experiments were approved by the institutional (QIMR Berghofer) ethics committee protocol A11605M.

#### Xenograft Transplantation Experiments

Primary AML samples were obtained from patients with AML, after informed consent in accordance with the Declaration of Helsinki, and approved by the institutional (QIMR Berghofer) ethics committee protocol P1382 (HREC/14/QRBW/278). Ficoll density gradient was then used to recover viable mononuclear cells. Viably frozen primary AML cells were thawed and CD3-depleted with biotinylated anti-human CD3 (SK7) and biotin-binder Dynabeads (Invitrogen) and subsequently injected via the lateral tail vein into 2.8 Gy irradiated (24h before transplant) NSG, NSGS or NRGS recipients. For normal hematopoiesis studies, viable mononuclear cells were isolated from cord blood samples by ficoll density gradient, CD3-depleted as above, and subsequently enriched for CD34+ cells using the human CD34 MicroBead kit (130-046-702 MACS Miltenyi Biotec) according to the manufacturers' instructions. 56,000 cells (donor 1) or 212,500 cells (donor 2) were injected via the lateral tail vein per irradiated NSG recipient.

#### In Vivo Liproxstatin-1 Treatment Studies

Liproxstatin-1 (SEL-S7699; Jomar Life Research) was dissolved in DMSO (7.9 mg in 400 μl), and then diluted with 2.44 ml 0.9% NaCl (Saline). Liproxstatin-1 (15 mg/kg i.p.) was administered by intraperitoneal (IP) injection via 27G insulin needle twice daily for 2 weeks (200 μl per recipient).

#### Blood Analysis

Blood was collected into EDTA-coated tubes and analyzed on a Hemavet 950 analyzer (Drew Scientific). PB smears were stained with Wright-Giemsa according to the manufacturer's protocol (BioScientific).

#### Histology

Tissues were fixed in 10% neutral buffered formalin, embedded in paraffin, and stained with H&E. Images of histological slides were obtained on a ScanScope FL (Aperio).

#### Flow Cytometry Analysis of AML PDX and Cord Blood Transplants

For monitoring human AML cell engraftment, 25-50 μl of PB was stained after red blood cell lysis (BD Pharmlyse, BD Biosciences) with anti-human CD45-AF647 (H130) and anti-mouse CD45.1-PE (A20). For AML phenotyping, cell populations were purified from BM (both femurs and tibiae) or SPL after red blood cell lysis and stained with anti-human CD45-FITC (H130), anti-mouse Cd45.1-PerCP/Cy5.5 (A20), anti-human CD34-PE (581), anti-human CD33-APC (WM53), anti-human CD38-APC/Cy7 (HIT2), and anti-human GPR56-PE/Cy7 (CG4). Flow cytometry analysis of lipid peroxidation was performed using C11-Bodipy 581/591 (Sapphire Bioscience) according to a previously published protocol (https://doi.org/10.1007/978-1-0716-0247-8_11), and reactive oxygen species were quantified using CellROX green (Invitrogen) according to the manufacturer's instructions, subsequent to cell surface marker staining. For the analysis of normal hematopoiesis, PB and SPL cells were stained after red blood cell lysis with anti-human CD45-FITC (H130), anti-mouse Cd45.1-PE (A20), anti-human CD19-PerCP/Cy5.5 (HIB19), anti-human CD33-APC (WM53), and anti-human CD3-APC/Cy7 (SK7). BM cells were isolated from both femurs and tibiae, red cell lysed, and stained with anti-human CD45-FITC (H130), anti-mouse Cd45.1-PerCP/Cy5.5 (A20), anti-human CD34-PE (581), anti-human CD38-APC/Cy7 (HIT2), and anti-CD33-APC (WM53). In all analyses, dead cells were discriminated by Sytox blue (Invitrogen). Washes and stainings were performed in PBS + 2% FCS + 1mM EDTA. Centrifugation steps were performed at 300 x g for 10 min at 4ºC. Flow cytometry analysis was performed on a FACS LSR Fortessa (BD Biosciences). Post-acquisition analyses were performed with FlowJo software V10.8.1 (Becton Dickinson & Company; BD).

**Cell Culture and In Vitro Cell-Growth Analysis**

AML cell lines were obtained from ATCC or DSMZ, Prof. Wallace Langdon (MOLM13, MV411, PL21), or Dr. S. Froehling (Monomac6; MM6). The identity of all cell lines was confirmed by STR profiling. Cells were cultured in RPMI supplemented with 10% fetal calf serum, 2 mM glutamine, and 200 U/ml penicillin, 200 μg/ml streptomycin. AML cells were seeded into a flat-bottom 96-well plate at a density of 2,500 cells per 100 ul final volume. Every 48-72 h, 25 or 50 ul of the cultures were transferred into a new plate, depending on the density of each cell line in the control condition, and supplemented with fresh medium containing imetelstat (GRN163L), mismatch controls (Mismatch 1 also referred to as MM1 or GRN140833; Mismatch 2 also referred to as GGG-mismatch, MM2 or GRN142865), or additional drugs of interest, i.e. ferrostatin-1 (Sigma-Aldrich), liproxstatin-1 (Sigma-Aldrich), desferrioxamine (DFOM; Hospira), zileuton (Sigma-Aldrich), menadione (Sigma-Aldrich), (+)-etomoxir sodium salt hydrate (Sigma-Aldrich), 1S,3R-RSL3 (Sigma-Aldrich), or erastin (Sigma-Aldrich). Cells were analyzed with Celltiter 96 aqueous nonradioactive cell proliferation assay (MTS Systems) according to the manufacturer’s instructions (Promega). Endpoint absorbance at 490 nm was detected using Biotek PowerWave and Gen5 data analysis software. Drug synergy scores were computed using the Synergyfinder 2.0 algorithm (https://synergyfinder.fimm.fi/).

**Flow cytometry analysis of AML cell lines**

Before staining, 2x10^5 cells were washed with phosphate-buffered saline (PBS) with 2% FCS and 1 mM EDTA. For cell cycle and G-quadruplex analysis, cells were then fixed and permeabilized using FIX & PERM Cell Permeabilization Kit (GAS-004; Invitrogen) and incubated with Anti-DNA G-quadruplex structures Antibody, clone BG4 (MABE917; Merck Millipore), and 0.2 mg/mL Hoechst 33342 (Invitrogen).

Flow cytometry analysis of lipid peroxidation was performed using C11-Bodipy 581/591 (Sapphire Bioscience) according to a previously published protocol (https://doi.org/10.1007/978-1-0716-0247-8_11), and reactive oxygen species were quantified using CellROX green (Invitrogen) according to the manufacturer's instructions. For both analyses, Sytox Blue 1.25 µM (S34857; Invitrogen) was used to distinguish between viable and dead cells. Flow cytometry analysis was performed on a FACS LSR Fortessa (BD Biosciences). Post-acquisition analyses were performed with FlowJo software V10.8.1 (Becton Dickinson & Company; BD).

**Imaging flow cytometry**

Lipophagy was detected by assessing co-localization of C12-BODIPY and LAMP-1 using a previously published method with modifications[3]. In detail, $1x10^6$ cells were harvested and washed in warm PBS (without FCS). Cells were then resuspended in warm RPMI (without FCS) containing 200 ng C12 FL Bodipy (Thermo Fisher Scientific) per ml, and incubated for 30 min at 37ºC. Cells were then washed in 9 ml wash buffer (PBS + 2% FCS + 1mM EDTA), and subsequently fixed and permeabilized using the FIX & PERM Cell Permeabilization Kit (GAS-004; Invitrogen) and incubated with Alexa Fluor® 647 anti-human CD107a (LAMP-1; Biolegend) according to the manufacturer's instructions. Cells were subsequently washed and resuspended in wash buffer with 0.2 mg/mL Hoechst 33342 (Invitrogen). Acqusition was performed using an Amnis® ImageStream®X Mark II Imaging

Flow and data were analyzed with IDEAS (Image Data Exploration and Analysis Software).

**Cloning guide RNAs into Plko5 vector**

Primers (Table below) were phosphorylated by using T4 polynucleotide kinase (EK0031; Thermo Scientific), T4 DNA ligase buffer (46300018; Invitrogen) and were incubated at 37°C for 45 minutes, inactivated at 70°C for 5 minutes and cooled down at 10°C. Plko5 vector was digested by using 10X Tango Yellow Buffer (BY5; Thermo Scientific), BsmBI (ER0451; Thermo Scientific) and 10 mM DTT (Thermo Scientific) and incubated at 37°C for 2 hours. FastAP Thermosensitive Alkaline Phosphatase (EF0654; Thermo Scientific) was added and vector was incubated at 37°C for 20 minutes, followed by 65°C for 20 minutes. Gel electrophoresis was performed and DNA was extracted by QIAQuick DNA Extraction Kit (QIAGEN). Then, 1:500 diluted, phosphorylated primers were ligated into digested Plko5 vector using T4 DNA ligase (New England BioLabs) and T4 DNA ligase buffer (New England BioLabs) by incubating at 22°C for 20 minutes and 65°C for 10 minutes. Vector without insert oligo was set up as a control. Vectors were transformed into E. coli and cultured on ampicillin plates and plasmid DNA was then purified by using Plasmid Maxi Kit (QIAGEN). Insertion of sgRNAs into Plko5 vector was confirmed by sequencing.

*Table: Single guide RNAs cloned into Plko5*

| Target gene | Forward primer | Reverse primer |
|---|---|---|
| ACSL4 sg1 | CACCGGTGTGTCTGAGGAGATAGCG | AAACCGCTATCTCCTCAGACACACC |
| ACSL4 sg2 | CACCGGCATCATCACTCCCTTAGGT | AAACACCTAAGGGAGTGATGATGCC |
| ACSL4 sg3 | CACCGACCTGGTCAGAGAGTGTAAG | AAACCTTACACTCTCTGACCAGGTC |
| ACSL4 sg4 | CACCGAAGCCCACTTCAGACAAACC | AAACGGTTTGTCTGAAGTGGGCTTC |
| FADS2 sg1 | CACCGTCTGGTACTGGAAATACATG | AAACCATGTATTTCCAGTACCAGAC |
| FADS2 sg2 | CACCGCCACGAATTCCAGGTCAGGG | AAACCCCTGACCTGGAATTCGTGGC |
| VIM sg1 | CACCGACAGCATGTCCAAATCGATG | AAACCATCGATTTGGACATGCTGTC |
| VIM sg2 | CACCGTCCGGTTGGCAGCCTCAGAG | AAACCTCTGAGGCTGCCAACCGGAC |
| VIM sg3 | CACCGTCTTGACCTTGAACGCAAAG | AAACCTTTGCGTTCAAGGTCAAGAC |
| VIM sg4 | CACCGCAACGACAAAGCCCGCGTCG | AAACCGACGCGGGCTTTGTCGTTGC |

*CD33 guide RNAs were used as described previously [4]*

*Table: Single guide RNA target sequences*

| Target gene | Target sequence | Strand | Exon |
|---|---|---|---|
| ACSL4 sg1 | ATCGGTGTGTCTGAGGAGATAGCGGGGCCC | Antisense | 12 |
| ACSL4 sg2 | TGATGCATCATCACTCCCTTAGGTCGGCCA | Antisense | 8 |
| ACSL4 sg3 | TGTCACCTGGTCAGAGAGTGTAAGCGGAGA | Antisense | 9 |
| ACSL4 sg4 | AGCTAAGCCCACTTCAGACAAACCTGGAAG | Sense | 4 |
| FADS2 sg1 | ATGATCTGGTACTGGAAATACATGGGGATG | Antisense | 7 |
| FADS2 sg2 | TTGCCCACGAATTCCAGGTCAGGGTGGAAG | Antisense | 2 |
| VIM sg1 | AGGAACAGCATGTCCAAATCGATGTGGATG | Sense | 5 |
| VIM sg2 | TTGTTCCGGTTGGCAGCCTCAGAGAGGTCA | Antisense | 6 |
| VIM sg3 | CACGTCTTGACCTTGAACGCAAAGTGGAAT | Sense | 4 |
| VIM sg4 | TAACCAACGACAAAGCCCGCGTCGAGGTGG | Sense | 2 |

PAM sequence highlighted in blue

### Transfection and Viral Titer Determination

Lentiviral particles were produced by transfection of 75% confluent HEK293T cells using Fugene (Promega) with pMD2.G (12559; Addgene) and psPAX2 (12260; Addgene) and Plko5 containing sgRNAs or empty backbone. Viral supernatants were harvested at 42 and 54 hours after transfection and filtered (45 nm pore size). Viral titer was determined by transduction of serial 1:10 dilutions of lentivirus into 3T3 cells using 1:1000 polybrene (Sigma Aldrich).

### Generation of Cas9-expressing Cell Lines

*Streptococcus pyogenes* Cas9 was stably introduced into the NB4, MV411, KO52 and TF1 cell lines by lentiviral infection. For Cas9/Blasticidin clones, a vector containing a human codon-optimised *Strep. Pyogenes* Cas9 and blasticidin resistance construct expressed from an EFS promoter (pFUGWb) was obtained from Addgene (lentiCas9-Blast, plasmid #52962) [5]. 0.5 x $10^6$ cells per well were seeded into a 6-well plate in a volume of 1.2 mL. 300 µL of concentrated virus was added to each well with polybrene at a final concentration of 8 µg/mL and centrifuged at 2,500 rpm for 90 minutes at 37°C. Cells were then incubated at 37°C for 48 hours with media replaced at 24 hours.

### Lentiviral Transduction and Sorting

NB4-, MV411-, KO52-, and TF1- expressing Cas9 cells were selected by incubation with blasticidin S HCl (Thermo Scientific; 5 µg/mL for NB4, MV411 and TF1; 1.25 µg/mL for KO52 cells) for two weeks. Cells were then transduced twice with lentivirus using 8 µg/mL polybrene (Sigma-Aldrich) and

0.01 M Hepes (Thermo Scientific), and were incubated overnight. Then, medium was refreshed and cells were incubated for 48 hours. Final multiplicity of infection (MOI) was 8. In order to select transduced cells, mCherry+ and Sytox blue- cells were sorted using a FACS AriaIIIu (BD Biosciences), when transduction efficiencies were <96%. CD33, VIM and mCherry expression were analyzed by flow cytometry after passage one. Cells were then stained with 0.6 µg/mL PerCPCy5.5 anti-human CD33 (303414; BioLegend) and 0.05 µM Sytox Blue (S34857; Invitrogen), or anti-human VIM (Vimentin (D21H3) XP® Rabbit mAb (Alexa Fluor® 488 Conjugate) #9854; Cell Signaling) after fixation and permeabilization using the FIX & PERM Cell Permeabilization Kit (GAS-004; Invitrogen). Flow cytometry analysis was performed on FACS LSR Fortessa (BD Biosciences) using DIVA software (BD Biosciences). Analyses were performed using FlowJo software V10.8.1 (Becton Dickinson & Company; BD).

#### Tracking of Indels by Decomposition

DNA was extracted using the DNeasy Blood and Tissue Kit (QIAGEN) and amplified using GoTaqGreen Master Mix (Promega, M7123).

PCR cycling conditions were: 2 minutes 95°C; 30 cycles of 30 seconds at 95°C, 30 seconds at 65°C and 90 seconds at 72°C; and a 5 minute final extension phase at 72°C. The PCR product was analyzed by agarose gel electrophoresis and extracted using the QIAquick gel extraction kit (QIAGEN, 28704) as per manufacturer's protocol. DNA was sequenced with a big dye terminator sequencer and editing efficiency was estimated using TIDE analysis (*https://tide.nki.nl/*).

*Table: Amplification of guide RNA target sequence contexts for TIDE analysis*

| Target gene | Forward primer | Reverse primer |
|---|---|---|
| FADS2 sg1 | CGCCGAGAGTTGCTCTGTTGCA | GGGGAAGGGAGATGGCCACACT |
| FADS2 sg2 | AGCCTGGTGCTGGCTCATCTCT | GCTCTGAGGCCCTCCCTGAACA |

*Table: PCR product sizes and sequencing primers for TIDE analysis*

| Target gene | PCR product size [bp] | Sequencing primer |
|---|---|---|
| FADS2 sg1 | 1242 | TGAGAGGCAGCTACCCCAGGGA |
| FADS2 sg2 | 1158 | GCTCTGAGGCCCTCCCTGAACA |

**Western Blotting**

Cells were lysed on ice using m-PER lysis buffer (Thermofisher scientific) supplemented with a protease and phosphatase inhibitor (cell signalling #5872) and protein was quantified using Pierce BCA protein assay kit (Thermofischer scientific). In total, 20-50 $\mu$g of protein extract was electrophoresed on a 4-15% SDS gradient gel (Mini-PROTEAN TGC Gel, Bio-rad) and transferred to an activated PVDF membrane for 1 h at 4 °C. Unspecific binding sites were blocked in 5% BSA in Tris-buffered saline with 1% Tween-20 (TBS-T) for 1 h at 4 °C. Primary antibodies were incubated overnight at 4 °C in constant motion in 5% BSA-TBS-T. The mouse anti-Cas9 antibody (#14697, Cell signalling), and rabbit anti-ACSL4 antibody (EPR8640, abcam) were used at 1:1,000 in 5% BSA-TBS-T. The mouse anti-ACTIN antibody (612656, BD biosciences) was used at 1:3,000 in 5% BSA-TBS-T. The membrane was washed with TBS-T 3 times for 5 minutes before incubation with the secondary antibody for 1 h at room temperature. Polyclonal goat anti-rabbit immunoglobulins HRP (P0448, Dako) and polyclonal rabbit anti-mouse (P0260, Dako) were used at 1:4,000. After subsequent membrane washing, protein was detected using Immobilon chemiluminescent HRP substrate (WBKLS0500, Millipore) and imaged with the iBright CL1500 imaging system.

**Terminal Restriction Fragment (TRF) Analysis**

Terminal restriction fragments were obtained from genomic DNA by complete digestion with the restriction enzymes HinfI and RsaI. TRFs were separated by pulsed-field gel electrophoresis. Gels were dried, denatured and subjected to in-gel hybridization with a γ-[32P]-ATP-labeled (CCCTAA)4 oligonucleotide probe. Gels were washed and the telomeric signal visualized by PhosphorImage analysis (Conomos et al., 2012). TRFs were processed by ImageJ 1.52a analysis software to quantitate mean telomere length.

**Telomere Length Q-PCR**

Samples were purified using the DNeasy Blood and Tissue Kit (QIAGEN). DNA isolation was performed as described previously [6], including degassing of buffers and supplementation with 50 μM of phenyl-tert-butyl nitrone to minimise oxidative damage. Telomere length was assessed using Q-PCR [7, 8]. A synthetic oligonucleotide of known length was serially diluted and utilized as a reference standard for telomere length and single copy gene to determine absolute telomere length (O'Callaghan 2011). Haemoglobin (HBB) and albumin were utilised as single-copy genes to calculate relative telomere/single copy gene (T/S) values and normalize telomere length per cell. Telomere length was validated by terminal restriction fragment (TRF) analysis, and four controls from TRF were included in each run to detect variance, including a long telomere control cell line.

The primers used for amplification were teloF, teloR [6], albu and albd [9] and HBB-f, 5' TGT GCT GGC CCA TCA CTT TG 3' and HBB-r 5' ACC AGC CAC CAC TTT CTG CTA GG 3'. For each sample 5-10 ng of sample DNA were loaded in triplicate. Primers were used at concentrations of 2 μM of forward primer and 18μM of reverse primer for telomere amplification and 6 μM and 14 μM respectively for single copy genes. DNA was amplified using Quantitect SYBR Green PCR Kit (QIAGEN) in a total volume of 20uL, and analyzed using an Applied Biosystem ViiA7 thermocycler.

**Brunello Genome-wide CRISPR Screen**

**Brunello Library**

The Brunello genome-wide gRNA library contains 76,441 gRNAs targeting 19,114 genes and was obtained from Addgene (Cat# 73178) [10]. Lentivirus containing the Brunello library was generated.

**Infection of Cells with Brunello guide RNA library for Pooled Screens**

Optimal infection conditions were determined for each batch of virus in the cell line of interest aiming to achieve 30-50% infection efficiency (corresponding to multiplicity of infection (MOI) of ~0.5-1). Infections of the Brunello library were performed in 12-well plates, seeding 3.0 x $10^6$ NB4 + Cas9 cells per well in a total volume of 2 mL. Cells were infected with 25 µL concentrated Brunello library virus with polybrene at a final concentration of 4 µg/mL by centrifuging plates at 2500 rpm for 90 minutes at 37°C. Infection efficiency was determined after 24 hours of incubation at 37°C.

To calculate infection efficiency, cells were replated in puromycin at 2 µg/mL selection dose. Infection efficiency was calculated after 48 hours in puromycin (at the time when non-infected controls were killed by puromycin) by comparing survival of cells with and without puromycin and calculated by the equation:

$$\left(\frac{\#\,Infected\ with\ Puro}{\#\,Infected\ without\ Puro} - \frac{\#\,Uninfected\ with\ Puro}{\#\,Uninfected\ without\ Puro}\right) \times 100$$
$$= Infection\ Rate\ (\%)$$

Cell populations ~30-50% infection efficiency underwent selection for gRNA-containing cells in 2 µg/mL of puromycin for 48 hours before being used for screening.

**DNA Preparation for guide RNA Sequencing**

Genomic DNA was extracted using Qiagen DNA kits (appropriate to cell number) according to the manufacturer's protocol. PCR of DNA was performed to attach Illumina sequencing adaptors and barcodes during the amplification of the gRNA region from each cell. Each 100 µL reaction contained a maximum of 10 µg of DNA plus $H_2O$ to a total volume of 50 µL, PCR master mix (40 µL) and 10 µL of a uniquely barcoded P7 primer (5 µM stock) allocated to each replicate. For each reaction, the 40 µL PCR master mix consisted of 0.75 µL of ExTaq DNA Polymerase (Clontech), 10 µL of 10x ExTaq buffer, 8 µL of deoxyribonucleotide triphosphate (dNTP) mix provided with the enzyme, 0.5 µL of P5 stagger primer mix (100 µM stock) and 20.75 µL $H_20$. The P5 primers attached to a common sequence of the gRNA vector 5' to the 20 nt gRNA insert with a stagger region integrated to increase diversity of the reads allowing the sequencer to recognize each read. P7 primers bound 3' to the gRNA insert and contain a unique 8 nt barcode to allow assignment of each read to a condition. PCR cycling conditions were: 1 minute 95°C; 28 cycles of 30 seconds at 94°C, 30 seconds at 52.5°C and 30 seconds at 72°C; and a 10 minute final extension phase at 72°C. The expected product size was 354 bp. P5/P7 primers were synthesized at IDT and are listed in the tables below.

**Table: Primer sequences for Brunello library Illumina Sequencing**

| Name | Sequence 5'-3' |
|---|---|
| P5 0 nt | AATGATACGGCGACCACCGAGATCTACACTCTTTCCCTACACGACGCTCTTCCGATCTTTGTGGAAAGGACGAAACACCG |
| P5 1 nt | AATGATACGGCGACCACCGAGATCTACACTCTTTCCCTACACGACGCTCTTCCGATCT**C**TTGTGGAAAGGACGAAACACCG |
| P5 2 nt | AATGATACGGCGACCACCGAGATCTACACTCTTTCCCTACACGACGCTCTTCCGATCT**GC**TTGTGGAAAGGACGAAACACCG |
| P5 3 nt | AATGATACGGCGACCACCGAGATCTACACTCTTTCCCTACACGACGCTCTTCCGATCT**AGC**TTGTGGAAAGGACGAAACACCG |
| P5 4 nt | AATGATACGGCGACCACCGAGATCTACACTCTTTCCCTACACGACGCTCTTCCGATCT**CAAC**TTGTGGAAAGGACGAAACACCG |
| P5 6 nt | AATGATACGGCGACCACCGAGATCTACACTCTTTCCCTACACGACGCTCTTCCGATCT**TGCACC**TTGTGGAAAGGACGAAACACCG |
| P5 7 nt | AATGATACGGCGACCACCGAGATCTACACTCTTTCCCTACACGACGCTCTTCCGATCT**ACGCAAC**TTGTGGAAAGGACGAAACACCG |
| P5 8 nt | AATGATACGGCGACCACCGAGATCTACACTCTTTCCCTACACGACGCTCTTCCGATCT**GAAGACCC**TTGTGGAAAGGACGAAACACCG |

Colour coding: P5 flow cell attachment sequence, Illumina sequencing primer, Stagger region, Vector primer binding sequence

**Table: P7 Primer Sequences for Brunello Library Illumina Sequencing**

| Name | Sequence 5'-3' |
|---|---|
| P7 A01 | CAAGCAGAAGACGGCATACGAGAT**CGGTTCAA**GTGACTGGAGTTCAGACGTGTGCTCTTCCGATCTTCTACTATTCTTTCCCCTGCACTGT |
| P7 A02 | CAAGCAGAAGACGGCATACGAGAT**GCTGGATT**GTGACTGGAGTTCAGACGTGTGCTCTTCCGATCTTCTACTATTCTTTCCCCTGCACTGT |
| P7 A03 | CAAGCAGAAGACGGCATACGAGAT**TAACTCGG**GTGACTGGAGTTCAGACGTGTGCTCTTCCGATCTTCTACTATTCTTTCCCCTGCACTGT |
| P7 A04 | CAAGCAGAAGACGGCATACGAGAT**TAACAGTT**GTGACTGGAGTTCAGACGTGTGCTCTTCCGATCTTCTACTATTCTTTCCCCTGCACTGT |
| P7 A05 | CAAGCAGAAGACGGCATACGAGAT**ATACTCAA**GTGACTGGAGTTCAGACGTGTGCTCTTCCGATCTTCTACTATTCTTTCCCCTGCACTGT |
| P7 A06 | CAAGCAGAAGACGGCATACGAGAT**GCTGAGAA**GTGACTGGAGTTCAGACGTGTGCTCTTCCGATCTTCTACTATTCTTTCCCCTGCACTGT |
| P7 A07 | CAAGCAGAAGACGGCATACGAGAT**ATTGGAGG**GTGACTGGAGTTCAGACGTGTGCTCTTCCGATCTTCTACTATTCTTTCCCCTGCACTGT |
| P7 A08 | CAAGCAGAAGACGGCATACGAGAT**TAGTCTAA**GTGACTGGAGTTCAGACGTGTGCTCTTCCGATCTTCTACTATTCTTTCCCCTGCACTGT |
| P7 A09 | CAAGCAGAAGACGGCATACGAGAT**CGGTGACC**GTGACTGGAGTTCAGACGTGTGCTCTTCCGATCTTCTACTATTCTTTCCCCTGCACTGT |
| P7 A10 | CAAGCAGAAGACGGCATACGAGAT**TACAGAGG**GTGACTGGAGTTCAGACGTGTGCTCTTCCGATCTTCTACTATTCTTTCCCCTGCACTGT |
| P7 A11 | CAAGCAGAAGACGGCATACGAGAT**ATTGTCAA**GTGACTGGAGTTCAGACGTGTGCTCTTCCGATCTTCTACTATTCTTTCCCCTGCACTGT |
| P7 A12 | CAAGCAGAAGACGGCATACGAGAT**TATGTCTT**GTGACTGGAGTTCAGACGTGTGCTCTTCCGATCTTCTACTATTCTTTCCCCTGCACTGT |

Colour coding: P7 flow cell attachment sequence, **Barcode**, Illumina sequencing primer, Vector primer binding sequence

Samples were purified with AMPure XP beads (Beckman Coulter, Cat# A63880) using the manufacturer's protocol for right-side selection (to remove residual genomic DNA) and left-side selection (to remove excess primers and dNTPs) and sequenced using the NextSeq 550 Illumina platform.

**Guide RNA Sequencing Analysis**

The sequenced gRNA insert from each read was mapped to a reference file of each gRNA in the library. The STARS and RIGER CRISPR screen analysis tools were applied to the sequencing results[10] with analysis based on log2-transformed numbers of gRNA reads in 'End' samples after drug treatment compared to 'Input' samples.

**RNA Sequencing: Library Preparation**

RNA was isolated from a maximum of 0.5 x $10^6$ cells using the Qiagen RNeasy Micro kit according to the manufacturer's instructions. Replicates were taken from at least 2 different experiments. RNA samples were quantitated using the Nanodrop Spectrophotometer (Thermo Fischer Scientific) and the Qubit Fluorometer using the Qubit RNA HS Assay Kit (Molecular Probes) according to the manufacturer's recommendations. RNA integrity was confirmed using the RNA 6000 PICO Kit (Agilent Technologies) with analysis using the Agilent 2100 Bioanalyser (Agilent Technologies). Total RNA (100 µg) was used for NGS and prepared according to the NEBNext Ultra II RNA Library Prep Kit for Illumina (New England Biolabs, NEB; Cat# E7770S). Briefly, following mRNA isolation, fragmentation and priming, first and second strand cDNA were synthesised. The double-stranded cDNA was purified, A-tailed and ligated to adaptors. Following purification, the adaptor ligated DNA was enriched by PCR, purified and assessed for quality using the High Sensitivity DNA Kit (Agilent) on the Agilent 2100 Bioanalyser. Libraries were sequenced using a High output, single-end, 75 cycle (version 2) sequencing kit on the Illumina Nextseq 550 platform.

**RNA Sequencing: Data Analysis**

Reads were trimmed for adapter sequences using Cutadapt (version 1.11) and aligned using Spliced Transcripts Alignment to a Reference (STAR) (version 2.5.2a) [11] to the GrCH37 assembly using the gene, transcript, and exon features of Ensembl (release 70) gene model. Expression was estimated using RNA-Seq by Expectation Maximization (RSEM) (version 1.2.30). Transcripts with zero read counts across all samples were removed prior to analysis. Normalization of read counts was performed by dividing by 1 million reads mapped to generate counts per million (CPM), followed by the trimmed mean of M-values (TMM) method from the edgeR package [12]. For the differential expression analysis, reads were filtered but not normalized, since edgeR performs normalization (library size and RNA composition) internally. For the differential expression (DE) analyses, the glmFit function was used to fit a negative binomial generalized log-linear model to the read counts for each transcript. Using the glmLRT function, we conducted transcript-wise likelihood ratio tests for each genotype comparison. Principal component analysis (PCA) was also performed on all DE transcripts with FDR <.05. Gene set enrichment analysis (GSEA) of transcriptomics data was performed using GSEA from the Broad Institute [13]. P-values were generated from 1000 gene set permutations, excluding gene sets with more than 3000 genes or less than 5 genes against custom made gene sets and the Broad Institute's Hallmark database.

**Mutational Sequencing**

Genomic alterations were profiled using the HemePACT assay (Integrated Mutation Profiling of Actionable Cancer Targets related to Hematological malignancies). This assay uses solution phase hybridization-based exon capture and massively parallel DNA sequencing to capture all protein-coding exons and select introns of 585 actionable cancer related genes. Samples were molecularly barcoded, to allow optimal cost efficiency during the capture process as well as at the sequencing step. 250ng of genomic DNA was used for library construction. Pools of 12 samples equimolarly mixed were sequenced at the Genomics Core Laboratory at MSKCC, in one lane of a Hiseq 2500, using the SBS chemistry for paired end 100/100 reads. The average coverage was greater than 400-fold, with a minimum of 99% of the targeted sequences covered 30 fold.

Reads were aligned to the reference human genome (hg19) using the Burrows-Wheeler Alignment tool [14]. Local realignment and quality score recalibration were conducted using the Genome Analysis Toolkit (GATK) according to GATK best practices [15].

Somatic alterations were identified (single-nucleotide variants, small insertions/deletions (indels), and copy number alterations). Single-nucleotide variants were identified using UnifiedGenotyper and mu Tect [16]. All samples were paired (AML/Normal) and candidate genomique alteration were reviewed manually in the Integrative Genomics Viewer [17].

**Lipidomics: Sample Preparation**

Cells were harvested and washed twice with cold PBS. Cell pellets of one Million cells were used. Eighteen samples were randomized and a blank negative control extraction was included. All steps were performed on ice. Cell pellets were re-suspended in 10 µL of pre-chilled milliQ water before adding 200 µL of ice-cold butanol/methanol (1:1) containing 10 mM ammonium formate and 50 µg/mL antioxidant 2,6-di-tert-butyl-4-methylphenol (BHT) to extract lipids. An aliquot of 10 µL of a 1/10-diluted SPLASH Lipidomix (Avanti, pn 330707) was spiked into each sample to assess sample preparation and retention time variation. Samples were incubated in a Thermomixer for 1 hr at 4°C and 850 rpm, followed by centrifugation for 15 min at 16,000 rcf (4°C). Supernatants were removed and dried down using a vacuum concentrator.
For LC/MS analysis, dried samples were re-suspended in 50 µL of ice-cold methanol/toluene (9:1, v/v) containing 100 ng/mL 12-[[(cyclohexylamino)carbonyl]amino]-dodecanoic acid (CUDA). The CUDA standard was used to flag autosampler inaccuracies. A sample pool was prepared by combining 5 µL of each sample.

**Lipidomics: Targeted LC/MS analysis**

Lipidomics was performed according to a method by Huynh and co-workers [18] with slight modifications. The LC/MS platform consisted of a 1290 Infinity II UHPLC coupled to a 6470 QQQ mass spectrometer via AJS ESI source (Agilent, Santa Clara, USA). The mass spectrometer was operated in positive ionization mode acquiring data in scheduled multiple reaction monitoring (MRM). Quadrupoles 1 and 2 were set to unit resolution. The MRM transition list contained 20 lipid classes and 593 lipid species (excluding internal standards CUDA and SPLASH Lipidomix) with the lipid naming convention used here also adopted from Huynh and co-workers [18].
Separation was performed on a Zorbax Eclipse Plus C18 RRHD (1.8 µm, 95 Å, 2.1 x 100 mm) analytical column connected to a 2.1 x 5 mm guard column of the same resin. The autosampler and column temperature were set to 4°C

and 60°C, respectively. Solvent system and gradient were used as described previously [18] and sample injection was 3 µL.
Source conditions were as follows: Gas temperature 175°C, gas flow 11 L/min, sheath gas temperature and flow at 250°C and 10 L/min, respectively, nebulizer 20 psi, fragmentor 135, capillary voltage at +4750 V, nozzle voltage was zero.
Prior to running samples, the originally published MRM transition list (20 lipid classes and 636 lipid species) was assessed for the presence/absence of lipid species in the current sample matrix using the sample pool. This was performed by splitting the transition list, running 3 scheduled MRM methods with a larger retention time window of 5 min. Following the analysis, the final scheduled MRM assay contained 609 transitions from 606 lipid species (including 13 ISTD lipid species and 16 transitions) with retention time windows of 1.5-4 min (depending on the lipid class and species, and a cycle time of 1 s resulting in a maximum of 177 concurrent MRMs and a minimum dwell time of 3.2 ms. All samples were run in a randomized order bracketed by a sample pool as a quality control.

**Lipidomics: Data Analysis**

Skyline-daily software [19] was used for lipid species assignment (precursor/product ion pairs and retention time) and chromatographic peak integration based on Huynh and co-workers [18]. An indexed retention time (iRT) calculator, which was generated using internal standards as well as lipid species assigned from the sample pool run, was employed for a retention time predictor to increase confidence of lipid assignment in subsequent samples. Peak picking was manually inspected using Skyline's retention times – replicate comparison pane and adjusted accordingly by comparing retention time and chromatographic peak profile to the sample pool QC run. Furthermore, all peaks were manually checked for correct integration.
The downstream data processing and visualization was carried out with R package lipidr [20]. Skyline transition results and a file with sample annotation and grouping were imported into lipidr. Raw data quality was assessed by plotting total lipid intensities of each sample as well as intensity and retention time distributions of internal standards across samples. Log2 transformation

and probabilistic quotient normalization (PQN) was performed prior to statistical analysis. Sample variation was investigated by principal component analysis (PCA). A lipid set enrichment analysis was performed by ranking fold changes, calculating enrichment scores and estimating the significance of enrichment using a permutation algorithm [20]. The enrichment results were plotted as boxplot or trend line.
