## Supplemental Table 1 for "Imetelstat-Mediated Alterations in Fatty Acid Metabolism To Induce Ferroptosis As Therapeutic Strategy for Acute Myeloid Leukemia"

**Table: Top 300 VMR**

| Ensembl.Gene.ID | Associated.Gene.Name | Rank |
|---|---|---|
| ENSG00000090382 | LYZ | 1 |
| ENSG00000005381 | MPO | 2 |
| ENSG00000019582 | CD74 | 3 |
| ENSG00000107317 | PTGDS | 4 |
| ENSG00000198804 | MT-CO1 | 5 |
| ENSG00000038427 | VCAN | 6 |
| ENSG00000197746 | PSAP | 7 |
| ENSG00000075624 | ACTB | 8 |
| ENSG00000030582 | GRN | 9 |
| ENSG00000172236 | TPSAB1 | 10 |
| ENSG00000122862 | SRGN | 11 |
| ENSG00000163220 | S100A9 | 12 |
| ENSG00000204287 | HLA-DRA | 13 |
| ENSG00000196126 | HLA-DRB1 | 14 |
| ENSG00000087086 | FTL | 15 |
| ENSG00000198886 | MT-ND4 | 16 |
| ENSG00000234745 | HLA-B | 17 |
| ENSG00000085265 | FCN1 | 18 |
| ENSG00000156508 | EEF1A1 | 19 |
| ENSG00000167996 | FTH1 | 20 |
| ENSG00000210082 | MIR4485 | 21 |
| ENSG00000251562 | MALAT1 | 22 |
| ENSG00000160255 | ITGB2 | 23 |
| ENSG00000124942 | AHNAK | 24 |
| ENSG00000231389 | HLA-DPA1 | 25 |
| ENSG00000198899 | MT-ATP6 | 26 |
| ENSG00000105223 | PLD3 | 27 |
| ENSG00000177575 | CD163 | 28 |
| ENSG00000117984 | CTSD | 29 |
| ENSG00000198727 | MT-CYB | 30 |
| ENSG00000140968 | IRF8 | 31 |
| ENSG00000157601 | MX1 | 32 |
| ENSG00000198938 | MT-CO3 | 33 |
| ENSG00000179348 | GATA2 | 34 |
| ENSG00000120738 | EGR1 | 35 |
| ENSG00000198786 | MT-ND5 | 36 |
| ENSG00000010327 | STAB1 | 37 |
| ENSG00000117289 | TXNIP | 38 |
| ENSG00000128342 | LIF | 39 |
| ENSG00000163131 | CTSS | 40 |
| ENSG00000101439 | CST3 | 41 |
| ENSG00000143384 | MCL1 | 42 |
| ENSG00000179344 | HLA-DQB1 | 43 |
| ENSG00000223865 | HLA-DPB1 | 44 |
| ENSG00000211459 | J01415.23 | 45 |
| ENSG00000170345 | FOS | 46 |

| | | |
|---|---|---|
| ENSG00000123384 | LRP1 | 47 |
| ENSG00000205542 | TMSB4X | 48 |
| ENSG00000026025 | VIM | 49 |
| ENSG00000140287 | HDC | 50 |
| ENSG00000227507 | LTB | 51 |
| ENSG00000166710 | B2M | 52 |
| ENSG00000198502 | HLA-DRB5 | 53 |
| ENSG00000178209 | PLEC | 54 |
| ENSG00000172232 | AZU1 | 55 |
| ENSG00000100292 | HMOX1 | 56 |
| ENSG00000198763 | MT-ND2 | 57 |
| ENSG00000142798 | HSPG2 | 58 |
| ENSG00000025708 | TYMP | 59 |
| ENSG00000198712 | MT-CO2 | 60 |
| ENSG00000143546 | S100A8 | 61 |
| ENSG00000185215 | TNFAIP2 | 62 |
| ENSG00000198888 | MT-ND1 | 63 |
| ENSG00000213366 | GSTM2 | 64 |
| ENSG00000216490 | IFI30 | 65 |
| ENSG00000107738 | C10orf54 | 66 |
| ENSG00000206503 | HLA-A | 67 |
| ENSG00000101347 | SAMHD1 | 68 |
| ENSG00000100316 | RPL3 | 69 |
| ENSG00000204525 | HLA-C | 70 |
| ENSG00000125740 | FOSB | 71 |
| ENSG00000005379 | BZRAP1 | 72 |
| ENSG00000149273 | RPS3 | 73 |
| ENSG00000170458 | CD14 | 74 |
| ENSG00000119535 | CSF3R | 75 |
| ENSG00000197766 | CFD | 76 |
| ENSG00000160883 | HK3 | 77 |
| ENSG00000167658 | EEF2 | 78 |
| ENSG00000196924 | FLNA | 79 |
| ENSG00000244734 | HBB | 80 |
| ENSG00000124766 | SOX4 | 81 |
| ENSG00000135218 | CD36 | 82 |
| ENSG00000136167 | LCP1 | 83 |
| ENSG00000197249 | SERPINA1 | 84 |
| ENSG00000137154 | RPS6 | 85 |
| ENSG00000120129 | DUSP1 | 86 |
| ENSG00000204592 | HLA-E | 87 |
| ENSG00000103187 | COTL1 | 88 |
| ENSG00000075426 | FOSL2 | 89 |
| ENSG00000197629 | MPEG1 | 90 |
| ENSG00000142541 | RPL13A | 91 |
| ENSG00000179388 | EGR3 | 92 |
| ENSG00000165168 | CYBB | 93 |
| ENSG00000084234 | APLP2 | 94 |

| | | |
|---|---|---|
| ENSG00000157514 | TSC22D3 | 95 |
| ENSG00000137801 | THBS1 | 96 |
| ENSG00000000938 | FGR | 97 |
| ENSG00000186407 | CD300E | 98 |
| ENSG00000142192 | APP | 99 |
| ENSG00000071575 | TRIB2 | 100 |
| ENSG00000158715 | SLC45A3 | 101 |
| ENSG00000204388 | HSPA1B | 102 |
| ENSG00000127951 | FGL2 | 103 |
| ENSG00000100448 | CTSG | 104 |
| ENSG00000093072 | CECR1 | 105 |
| ENSG00000157404 | KIT | 106 |
| ENSG00000174059 | CD34 | 107 |
| ENSG00000254772 | EEF1G | 108 |
| ENSG00000129226 | CD68 | 109 |
| ENSG00000185507 | IRF7 | 110 |
| ENSG00000096060 | FKBP5 | 111 |
| ENSG00000070756 | PABPC1 | 112 |
| ENSG00000135318 | NT5E | 113 |
| ENSG00000128016 | ZFP36 | 114 |
| ENSG00000130066 | SAT1 | 115 |
| ENSG00000171223 | JUNB | 116 |
| ENSG00000167526 | RPL13 | 117 |
| ENSG00000160789 | LMNA | 118 |
| ENSG00000168685 | IL7R | 119 |
| ENSG00000056736 | IL17RB | 120 |
| ENSG00000137193 | PIM1 | 121 |
| ENSG00000155659 | VSIG4 | 122 |
| ENSG00000041353 | RAB27B | 123 |
| ENSG00000162511 | LAPTM5 | 124 |
| ENSG00000018280 | SLC11A1 | 125 |
| ENSG00000164733 | CTSB | 126 |
| ENSG00000115306 | SPTBN1 | 127 |
| ENSG00000196415 | PRTN3 | 128 |
| ENSG00000067225 | PKM | 129 |
| ENSG00000089157 | RPLP0 | 130 |
| ENSG00000132475 | H3F3B | 131 |
| ENSG00000269404 | SPIB | 132 |
| ENSG00000133112 | TPT1 | 133 |
| ENSG00000121316 | PLBD1 | 134 |
| ENSG00000137959 | IFI44L | 135 |
| ENSG00000184009 | ACTG1 | 136 |
| ENSG00000245532 | NEAT1 | 137 |
| ENSG00000158517 | NCF1 | 138 |
| ENSG00000140678 | ITGAX | 139 |
| ENSG00000140988 | RPS2 | 140 |
| ENSG00000152518 | ZFP36L2 | 141 |
| ENSG00000127528 | KLF2 | 142 |

| | | |
|---|---|---|
| ENSG00000177663 | IL17RA | 143 |
| ENSG00000198695 | MT-ND6 | 144 |
| ENSG00000028137 | TNFRSF1B | 145 |
| ENSG00000173546 | CSPG4 | 146 |
| ENSG00000124882 | EREG | 147 |
| ENSG00000148154 | UGCG | 148 |
| ENSG00000150991 | UBC | 149 |
| ENSG00000165646 | SLC18A2 | 150 |
| ENSG00000144668 | ITGA9 | 151 |
| ENSG00000067082 | KLF6 | 152 |
| ENSG00000145425 | RPS3A | 153 |
| ENSG00000160593 | AMICA1 | 154 |
| ENSG00000215252 | GOLGA8B | 155 |
| ENSG00000147604 | RPL7 | 156 |
| ENSG00000167470 | MIDN | 157 |
| ENSG00000111640 | GAPDH | 158 |
| ENSG00000231486 | AC096579.7 | 159 |
| ENSG00000163931 | TKT | 160 |
| ENSG00000078804 | TP53INP2 | 161 |
| ENSG00000175061 | C17orf76-AS1 | 162 |
| ENSG00000188404 | SELL | 163 |
| ENSG00000169385 | RNASE2 | 164 |
| ENSG00000105426 | PTPRS | 165 |
| ENSG00000251322 | SHANK3 | 166 |
| ENSG00000018510 | AGPS | 167 |
| ENSG00000105835 | NAMPT | 168 |
| ENSG00000204103 | MAFB | 169 |
| ENSG00000169896 | ITGAM | 170 |
| ENSG00000130592 | LSP1 | 171 |
| ENSG00000138722 | MMRN1 | 172 |
| ENSG00000026508 | CD44 | 173 |
| ENSG00000121064 | SCPEP1 | 174 |
| ENSG00000106484 | MEST | 175 |
| ENSG00000180530 | NRIP1 | 176 |
| ENSG00000178695 | KCTD12 | 177 |
| ENSG00000105472 | CLEC11A | 178 |
| ENSG00000136997 | MYC | 179 |
| ENSG00000175265 | GOLGA8A | 180 |
| ENSG00000125810 | CD93 | 181 |
| ENSG00000179583 | CIITA | 182 |
| ENSG00000134202 | GSTM3 | 183 |
| ENSG00000148303 | RPL7A | 184 |
| ENSG00000167880 | EVPL | 185 |
| ENSG00000211899 | IGHM | 186 |
| ENSG00000140718 | FTO | 187 |
| ENSG00000178573 | MAF | 188 |
| ENSG00000137818 | RPLP1 | 189 |
| ENSG00000184371 | CSF1 | 190 |

| | | |
|---|---|---|
| ENSG00000139318 | DUSP6 | 191 |
| ENSG00000146192 | FGD2 | 192 |
| ENSG00000131016 | AKAP12 | 193 |
| ENSG00000198034 | RPS4X | 194 |
| ENSG00000101160 | CTSZ | 195 |
| ENSG00000108551 | RASD1 | 196 |
| ENSG00000100097 | LGALS1 | 197 |
| ENSG00000131401 | NAPSB | 198 |
| ENSG00000106066 | CPVL | 199 |
| ENSG00000121053 | EPX | 200 |
| ENSG00000100368 | CSF2RB | 201 |
| ENSG00000008516 | MMP25 | 202 |
| ENSG00000177606 | JUN | 203 |
| ENSG00000166963 | MAP1A | 204 |
| ENSG00000128815 | WDFY4 | 205 |
| ENSG00000130429 | ARPC1B | 206 |
| ENSG00000160796 | NBEAL2 | 207 |
| ENSG00000100345 | MYH9 | 208 |
| ENSG00000142347 | MYO1F | 209 |
| ENSG00000130522 | JUND | 210 |
| ENSG00000120833 | SOCS2 | 211 |
| ENSG00000164929 | BAALC | 212 |
| ENSG00000162104 | ADCY9 | 213 |
| ENSG00000168209 | DDIT4 | 214 |
| ENSG00000120708 | TGFBI | 215 |
| ENSG00000105372 | RPS19 | 216 |
| ENSG00000104870 | FCGRT | 217 |
| ENSG00000198840 | MT-ND3 | 218 |
| ENSG00000111335 | OAS2 | 219 |
| ENSG00000135486 | HNRNPA1 | 220 |
| ENSG00000196628 | TCF4 | 221 |
| ENSG00000163563 | MNDA | 222 |
| ENSG00000159840 | ZYX | 223 |
| ENSG00000156675 | RAB11FIP1 | 224 |
| ENSG00000121691 | CAT | 225 |
| ENSG00000171282 | BAHCC1 | 226 |
| ENSG00000105640 | RPL18A | 227 |
| ENSG00000170571 | EMB | 228 |
| ENSG00000131042 | LILRB2 | 229 |
| ENSG00000132744 | ACY3 | 230 |
| ENSG00000248527 | RP5-857K21.10 | 231 |
| ENSG00000185198 | PRSS57 | 232 |
| ENSG00000196205 | EEF1A1P5 | 233 |
| ENSG00000228253 | MT-ATP8 | 234 |
| ENSG00000078596 | ITM2A | 235 |
| ENSG00000172247 | C1QTNF4 | 236 |
| ENSG00000183087 | GAS6 | 237 |
| ENSG00000171314 | PGAM1 | 238 |

| | | |
|---|---|---|
| ENSG00000112773 | FAM46A | 239 |
| ENSG00000052749 | RRP12 | 240 |
| ENSG00000180044 | C3orf80 | 241 |
| ENSG00000179820 | MYADM | 242 |
| ENSG00000092964 | DPYSL2 | 243 |
| ENSG00000169403 | PTAFR | 244 |
| ENSG00000142937 | RPS8 | 245 |
| ENSG00000168329 | CX3CR1 | 246 |
| ENSG00000233276 | GPX1 | 247 |
| ENSG00000114554 | PLXNA1 | 248 |
| ENSG00000203812 | HIST2H2AA4 | 249 |
| ENSG00000118971 | CCND2 | 250 |
| ENSG00000151726 | ACSL1 | 251 |
| ENSG00000083845 | RPS5 | 252 |
| ENSG00000231500 | RPS18 | 253 |
| ENSG00000126759 | CFP | 254 |
| ENSG00000110719 | TCIRG1 | 255 |
| ENSG00000138623 | SEMA7A | 256 |
| ENSG00000175899 | A2M | 257 |
| ENSG00000166278 | C2 | 258 |
| ENSG00000100906 | NFKBIA | 259 |
| ENSG00000106546 | AHR | 260 |
| ENSG00000096384 | HSP90AB1 | 261 |
| ENSG00000140105 | WARS | 262 |
| ENSG00000002586 | CD99 | 263 |
| ENSG00000105492 | SIGLEC6 | 264 |
| ENSG00000104951 | IL4I1 | 265 |
| ENSG00000206172 | HBA1 | 266 |
| ENSG00000111913 | FAM65B | 267 |
| ENSG00000126709 | IFI6 | 268 |
| ENSG00000010610 | CD4 | 269 |
| ENSG00000113140 | SPARC | 270 |
| ENSG00000141026 | MED9 | 271 |
| ENSG00000174444 | RPL4 | 272 |
| ENSG00000110324 | IL10RA | 273 |
| ENSG00000136068 | FLNB | 274 |
| ENSG00000138316 | ADAMTS14 | 275 |
| ENSG00000137076 | TLN1 | 276 |
| ENSG00000172216 | CEBPB | 277 |
| ENSG00000042493 | CAPG | 278 |
| ENSG00000116701 | NCF2 | 279 |
| ENSG00000074800 | ENO1 | 280 |
| ENSG00000184916 | JAG2 | 281 |
| ENSG00000104267 | CA2 | 282 |
| ENSG00000132530 | XAF1 | 283 |
| ENSG00000123358 | NR4A1 | 284 |
| ENSG00000183688 | FAM101B | 285 |
| ENSG00000177989 | ODF3B | 286 |

| | | |
|---|---|---|
| ENSG00000185686 | PRAME | 287 |
| ENSG00000189060 | H1F0 | 288 |
| ENSG00000091592 | NLRP1 | 289 |
| ENSG00000160932 | LY6E | 290 |
| ENSG00000135916 | ITM2C | 291 |
| ENSG00000155090 | KLF10 | 292 |
| ENSG00000197958 | RPL12 | 293 |
| ENSG00000008988 | RPS20 | 294 |
| ENSG00000108518 | PFN1 | 295 |
| ENSG00000198400 | NTRK1 | 296 |
| ENSG00000108679 | LGALS3BP | 297 |
| ENSG00000121966 | CXCR4 | 298 |
| ENSG00000182718 | ANXA2 | 299 |
